# Making the effect visible – OX40 targeting nanobodies for *in vivo* imaging of activated T cells

**DOI:** 10.1101/2024.08.09.607337

**Authors:** Desiree I. Frecot, Simone Blaess, Teresa R. Wagner, Philipp D. Kaiser, Bjoern Traenkle, Madeleine Fandrich, Meike Jakobi, Armin M. Scholz, Stefan Nueske, Nicole Schneiderhan-Marra, Cécile Gouttefangeas, Manfred Kneilling, Bernd J. Pichler, Dominik Sonanini, Ulrich Rothbauer

## Abstract

**Purpose:** Human OX40 (hOX40/CD134), a member of the TNF receptor superfamily, is mainly expressed on activated T lymphocytes. Triggered by its ligand OX40L (CD252), it provides costimulatory signals that support the differentiation, proliferation and long-term survival of T cells. Besides being a relevant therapeutic target, hOX40 is also an important biomarker for monitoring the presence or infiltration of activated T cells within the tumor microenvironment (TME), the inflammatory microenvironment (IME) in immune-mediated diseases (IMIDs) and the lymphatic organs. Here, we developed novel single domain antibodies (nanobodies, Nbs) targeting hOX40 to monitor the activation status of T cells by *in vivo* molecular imaging.

**Methods:** Nbs against hOX40 (hOX40-Nbs) were selected from an immunized Nb-library by phage display. The identified hOX40-Nbs were characterized *in vitro*, including determination of their specificity, affinity, stability, epitope recognition and their impact on OX40 signaling and T cell function. A lead candidate was site-specifically conjugated with a fluorophore via sortagging and applied for noninvasive *in vivo* optical imaging (OI) of hOX40-expressing cells in a xenograft mouse model.

**Results:** Our selection campaign revealed four unique Nbs that exhibit strong binding affinities and high stabilities under physiological conditions. Epitope binning and domain mapping indicated the targeting of at least two different epitopes on hOX40. When analyzing their impact on OX40 signaling, an agonistic effect was excluded for all validated Nbs. Incubation of activated T cells with hOX40-Nbs did not affect cell viability or proliferation patterns, whereas differences in cytokine release were observed. *In vivo* OI with a fluorophore-conjugated lead candidate in experimental mice with hOX40-expressing xenografts demonstrated its specificity and functionality as an imaging probe.

**Conclusion:** Considering the need for advanced probes for noninvasive *in vivo* monitoring of T cell activation dynamics, we propose, that our hOX40-Nbs have a great potential as imaging probes for noninvasive and longitudinal *in vivo* diagnostics. Quantification of OX40^+^ T cells in TME or IME will provide crucial insights into the activation state of infiltrating T cells, offering a valuable biomarker for assessing immune responses, predicting treatment efficacy, and guiding personalized immunotherapy strategies in patients with cancer or IMIDs.

## Introduction

Immunotherapies that specifically modulate the patient’s immune system, e.g. to fight malignant tumor cells or attenuate autoimmune reactions, have opened a new chapter in personalized medicine (1–4). Although such therapies have shown remarkable success in some cases, the reasons why patients respond differently need to be understood. It is generally accepted that treatment outcomes are highly dependent on the individual immune system and the composition of the tumor (TME) or inflammatory microenvironment (IME), which is why sophisticated diagnostic approaches are required. To overcome the limitations of invasive procedures including histopathology or liquid biopsies (5, 6), noninvasive techniques such as *in vivo* imaging have been implemented in diagnostics and therapy monitoring. This has led to an increasing interest in the development of novel probes, that recognize specific immune cell populations and are capable of visualizing their infiltration into the TME, IME as well as primary and secondary lymphatic organs (7). Beginning with CD8^+^ T cells as one of the most important immune cells in the context of immunotherapies (8, 9), preclinical and clinical *in vivo* imaging of other immune cells such as CD4^+^ T cells or myeloid cells including tumor-associated macrophages has been reported more recently (7, 10–13). Since the presence or absence of specific cell populations alone does not give information about their functional states, visualizing of particularly activation, could enable more precise patient stratification and monitoring of therapeutic responses. This was demonstrated for the inducible T cell costimulatory receptor (ICOS) (14, 15), the early T cell activation marker CD69 (16), and granzyme B, which is released by activated cytotoxic T cells or interferon-γ (IFN-γ) (17).

OX40 (CD134/ TNFRSF4), has been described as a surface marker for T cell activation (17–21). It is mainly expressed on activated CD8^+^ and CD4^+^ T cells, but also on activated regulatory T cells (Tregs), natural killer T cells (NKTs) and neutrophils (19, 22–24). OX40 binds to the OX40 ligand (OX40L, CD252) presented by activated antigen-presenting cells (APCs) including B cells, dendritic cells and macrophages (25–27). OX40-OX40L engagement is key to potentiate T cell responses, including differentiation, proliferation, long-term survival, and enhancement of T cell effector functions, such as cytokine production (28). Recently, mouse-specific OX40 monoclonal antibodies (mAbs) were developed and applied for immune positron emission tomography (immunoPET) imaging in proof-of-principle studies to predict responses to cancer vaccines (29) or T cell response to glioblastoma (30) in preclinical mouse models. Beside cancer, OX40-specific immunoPET has also been applied to follow the development of acute graft-versus-host disease (31) or rheumatoid arthritis (32). However, the long systemic half-life of mAbs (up to seven days after injection) delays prompt immunoPET imaging time points due to the associated high background signal and leads to high radiation exposure. Moreover, mAb-based immunoPET yields a high tissue and blood background, which limits its sensitivity in detecting small populations of activated T cells (33). Consequently, there is a high demand for advanced molecules targeting human OX40 (hOX40) for diagnostic immunoPET imaging (34).

Antibody fragments derived from heavy-chain-only antibodies of camelids, referred to as VHHs or nanobodies (Nbs) (35), have emerged as versatile medical *in vivo* imaging probes (reviewed in (36–38)). In combination with highly sensitive and/or quantitative whole-body molecular imaging techniques such as optical or radionuclide-based modalities, Nbs have been shown to bind their targets within several minutes of systemic application (37, 39). Here, we describe the first set of hOX40-specific Nbs to monitor the activation status of human T cells and demonstrate the capability of a lead candidate for whole-body *in vivo* optical imaging (OI) of hOX40-expressing tumor cells in a mouse xenograft model.

## Results

### Identification and characterization of hOX40 specific Nbs

For the generation of hOX40-specific Nbs, two alpacas (*Vicugna pacos*) were immunized with the extracellular domain of hOX40, which contains the amino acid residues from Leu29 to Ala216. A positive immune response in both animals was confirmed on day 63 by serum ELISA (**Supplementary Figure S1**). Starting from peripheral blood lymphocytes (PBLs), we established a Nb-phagemid library representing the VHH repertoire of both animals (size: ∼3 x 10^7^ clones), from which hOX40-Nbs were enriched against recombinant hOX40 in two consecutive rounds of phage display. The selective binding of individual clones was tested in a whole-cell phage ELISA with U2OS cells stably expressing hOX40 (U2OS-hOX40). Subsequent sequencing revealed four unique hOX40-Nbs, namely O7, O12, O18 and O19, which exhibited highly diverse complementarity determining regions (CDRs) 3 (**Figure 1A, Supplementary Table S1**). All selected Nbs were expressed in *Escherichia coli* (*E. coli)* and purified by immobilized metal ion affinity chromatography (IMAC) followed by size exclusion chromatography (SEC), yielding high purity binding molecules (**Figure 1B**). To initially assess their binding affinities, we performed biolayer interferometry (BLI) and determined K_D_ values in the pico-to low nanomolar range (0.2 - 3.4 nM), while O7 showed a substantially weaker affinity as reflected by a K_D_ of ∼ 140 nM (**Figure 1C, Supplementary Figure S2A**). In addition, we determined the folding stability of the selected candidates using differential scanning fluorimetry (nanoDSF). All Nbs showed high thermal stabilities with melting temperatures (T_M_) between 50°C and 74°C without aggregation. Notably, this was not affected by an accelerated aging period of 10 days at 37°C (**Figure 1D, Supplementary Figure S2B**).

**Fig. 1:**
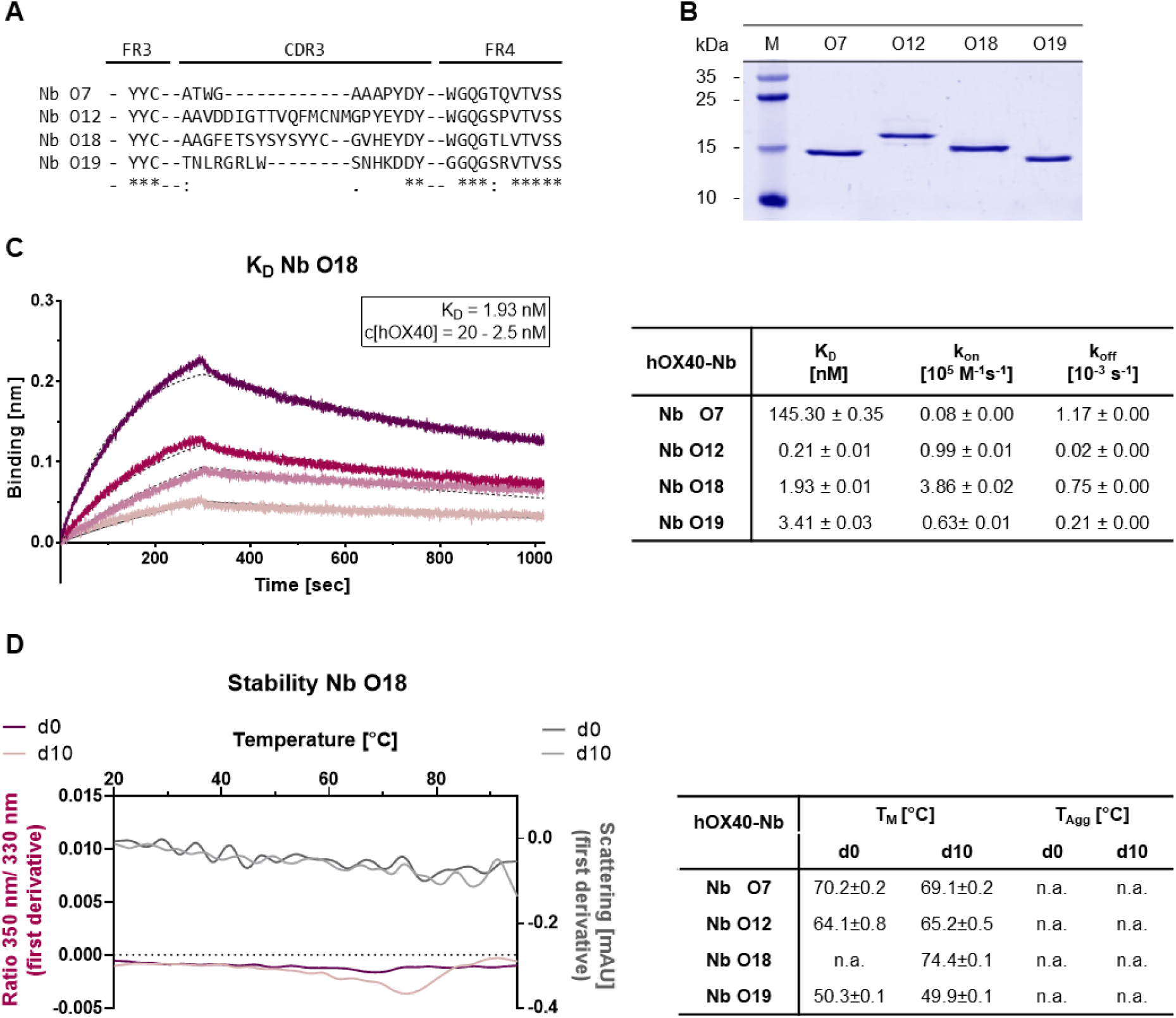
Biochemical characterization of hOX40-Nbs. (**A**) Amino acid (aa) sequences of the complementarity determining region (CDR) 3 from 4 unique hOX40-Nbs identified by two consecutive rounds of bio panning (full sequences are displayed in **Supplementary Table S1**). (**B**) Coomassie-stained SDS-PAGE of 2 µg purified hOX40-Nbs after purification using immobilized metal affinity chromatography (IMAC) and size exclusion chromatography (SEC). (**C**) Biolayer interferometry (BLI)-based affinity measurements exemplarily shown for Nb O18. Biotinylated Nb was immobilized on streptavidin biosensors. Kinetic measurements were performed using four concentrations of recombinant hOX40 ranging from 2.5 nM - 20 nM (displayed with gradually lighter shades of color; left). Summary table (right) shows affinities (KD), association constants (kon), and dissociation constants (koff) determined by BLI using four concentrations of purified Nbs as mean ± SD. (**D**) Stability analysis using nano scale differential scanning fluorimetry (nanoDSF) displaying fluorescence ratio (350 nm/330 nm) (red) and light scattering (gray) shown as first derivative for day 0 (dark shade) and after an accelerated aging period of 10 days at 37°C (light shade), exemplary shown for Nb O18 (left) and summarized for all hOX40-Nbs in the table (right). Data are shown as mean value of three technical replicates.

For the fluorescent functionalization of hOX40-Nbs, we took advantage of a sortase-based approach to selectively attach an azide-group to the C-termini of the Nbs, which served as chemical handle for the addition of a AlexaFluor647(AF647)-conjugated dibenzocyclooctyne (DBCO-AF647) group utilizing click chemistry (11). As a result, we obtained Nbs comprising a C-terminal fluorophore with a defined labeling ratio of 1:1. The fluorescent Nbs were used to determine corresponding EC_50_ values on U2OS-hOX40 cells by flow cytometry. In accordance with the BLI-determined affinities, a strong functional binding for O12 and O18 with EC_50_ values in the subnanomolar range (∼ 0.1 nM for O12; ∼ 0.3 nM for O18) was determined, whereas O7 and O19 displayed slightly weaker affinities (**Figure 2A, Supplementary Figure S3A**). To further confirm specific binding of the selected Nbs to hOX40 localized at the plasma membrane of mammalian cells, we used the fluorescent Nbs for live cell staining of U2OS-hOX40 cells in comparison to wild-type U2OS (U2OS-WT) cells. The images displayed intense signals localized at the cellular surface for all tested binders, which was comparable to the staining with a commercially available anti-hOX40 mAb, while none of the tested Nbs showed non-specific binding to U2OS-WT cells (**Figure 2B**). In addition, we used this approach to test a potential cross-reactivity of the Nbs to murine OX40 (mOX40) and performed live cell imaging on U2OS cells transiently expressing the mOX40. Only O7 bound to mOX40, while all other candidates showed no staining of mOX40 expressing U2OS cells (**Supplementary Figure S3B**). In summary, we identified four hOX40-Nbs that bind recombinant as well as cell-resident hOX40. With respect to O12, O18 and O19 we selected high-affinity binders with K_D_ values in the pico to low nanomolar range that exhibited strong specific binding to membrane-exposed hOX40. Notably, only O7 was less affine to recombinant hOX40 but showed additional cross-reactivity towards mOX40.

**Fig. 2:**
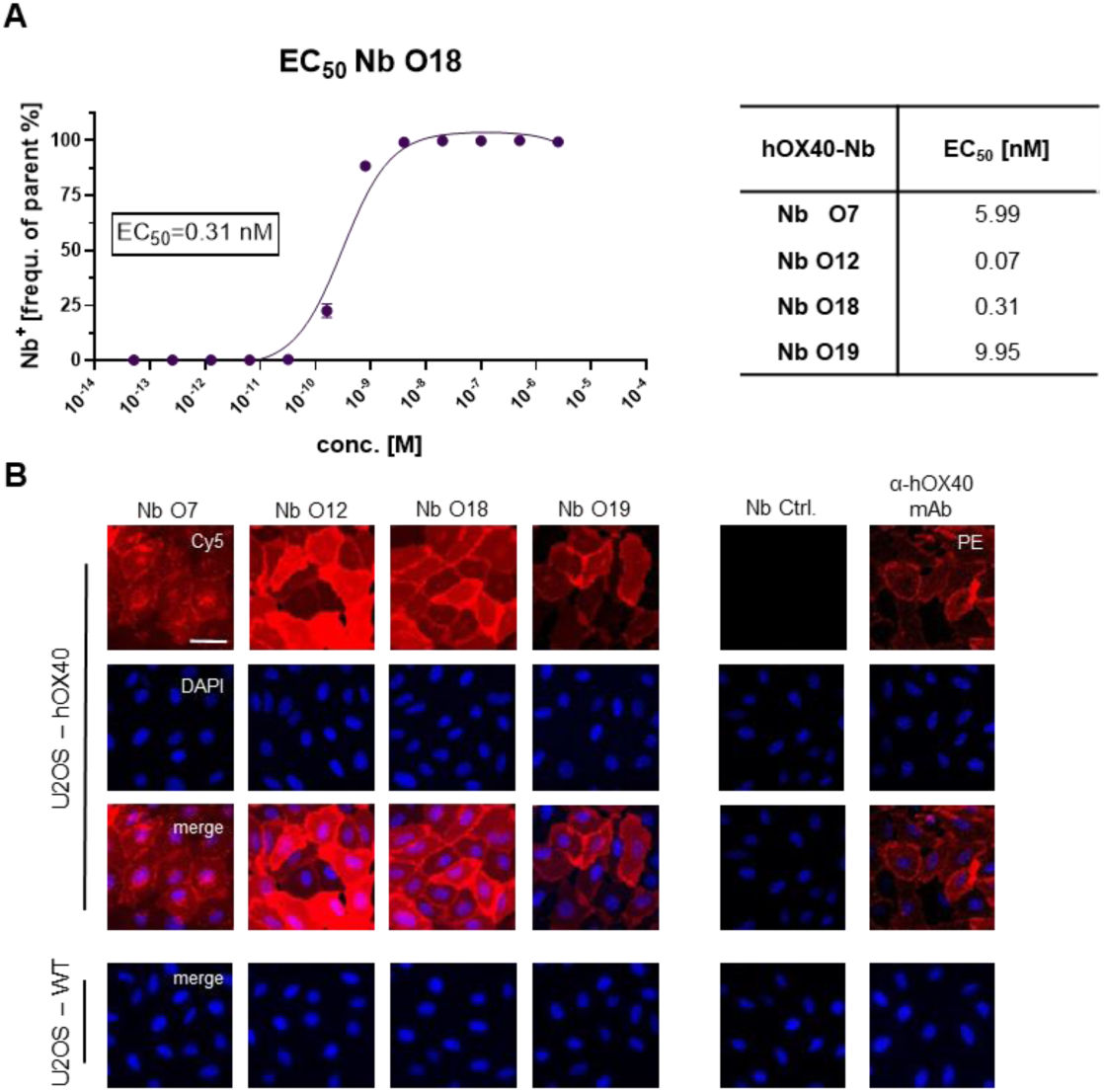
Characterization of cellular binding of hOX40-Nbs. (**A**) Determination of hOX40-Nb binding to cellular expressed hOX40 by flow cytometry (n=3), exemplary shown for Nb O18 labeled with AlexaFluor647 (AF647; left). The percentage of positively stained U2OS-hOX40 (frequency of parent) was plotted against indicated concentrations of AF647-labeled hOX40-Nbs and EC50 values shown in table (right) were calculated from a four-parametric sigmoidal model based on the mean ± SD of three technical replicates. (**B**) Representative images of U2OS-hOX40 cells (upper panel) and U2OS-WT cells (lower panel) stained with AF647-labeled hOX40-Nbs (left) as well as an unspecific Nb (Nb Ctrl.) as negative and phycoerythrin (PE)-labeled αnti-hOX40 mAb as positive control (right). Shown are individual Nb staining, nuclei staining (Hoechst, blue) and merged signals; scale bar: 50 µm.

### Characterization of binding epitopes on hOX40

To localize the binding sites of the selected hOX40-Nbs within the natively folded hOX40, we generated cellular expression constructs comprising domain-deletion mutants of hOX40 domains 1-3, which we transiently expressed in U2OS cells. Nb binding to truncated versions of hOX40 was visualized by immunofluorescence imaging of live cells. An anti-hOX40 mAb directed against domain 4 was used as a positive control (**Figure 3A**). Based on these results, we allocated binding of O7 and O19 to domain 3 and of O12 and O18 to domain 1 of hOX40 (**Figure 3B**). To examine a potential combinatorial binding of the different hOX40-Nbs, we further performed epitope binning analysis by BLI (**Figure 3C**). As expected from the domain mapping, O12 and O18, both targeting domain 1, simultaneously bound hOX40 in complex with O7 or O19, each targeting domain 3. However, only weak combinatorial binding was observed for Nbs addressing the same domain, suggesting that O12 and O18, as well as O7 and O19 address identical or at least overlapping epitopes (**Figure 3D, Supplementary Figure S4**).

**Fig. 3:**
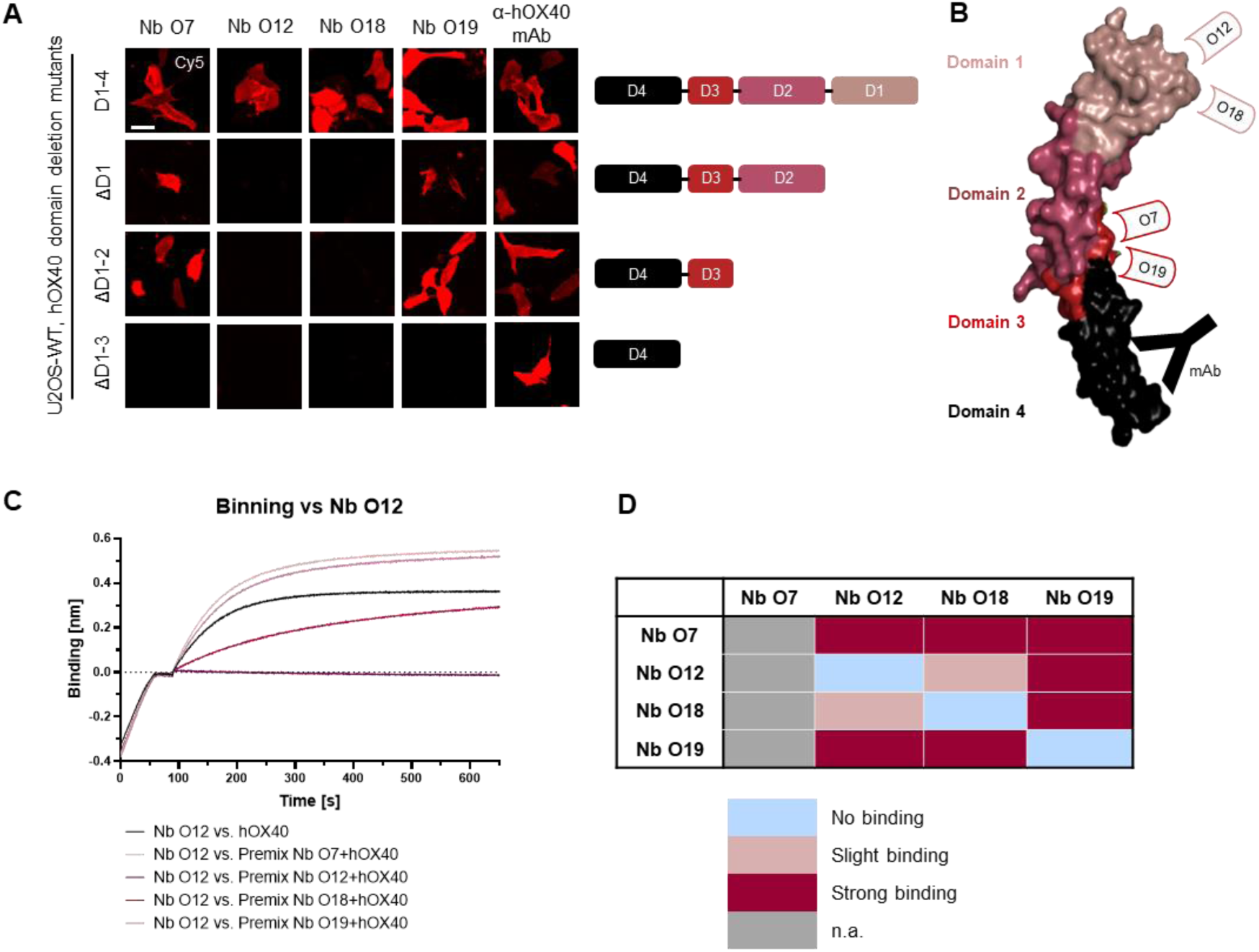
Characterization of binding epitopes of hOX40-Nbs. (**A**) Domain mapping by immunofluorescence staining with hOX40-Nbs on U2OS cells displaying either surface exposed hOX40 full length (D1-4), or domain deletion mutants as indicated. Shown are representative images of living cells stained with individual AF647-labeled Nbs or anti-hOX40 mAb; scale bar: 50 µm. (**B**) Schematic overview summarizing the results of domain mapping analysis (crystal structure OX40 PDB: 2HEV). (**C**) Epitope binning analysis of hOX40-Nbs by BLI. Representative sensograms of combinatorial Nb binding to recombinant hOX40 on sharing/overlapping epitopes or on different epitopes are shown (left). (**D**) Graphical summary of epitope binning analysis.

### hOX40-Nbs bind to activated human T lymphocytes

Having demonstrated that all selected Nbs recognize recombinant and exogenously overexpressed cellular hOX40, we next investigated their specificity for binding to endogenous hOX40 on activated T cells. Therefore, human peripheral blood mononuclear cells (hPBMCs) from three healthy donors (K025, K029 and K034) were either left untreated or incubated for 24 h with phytohaemagglutinin L (PHA-L) as a pan T cell stimulus to induce expression of OX40 (40) (**Figure 4A**). Subsequently, hPBMCs were double-stained with fluorescently labeled hOX40-Nbs or a phycoerythrin (PE)-labeled anti-hOX40 mAb in combination with a T cell-specific anti-CD3 mAb, and the percentage of double-positive T cells (CD3^+^OX40^+^) was analyzed by flow cytometry (**Figure 4C**, gating: **Supplementary Figure S5**). All hOX40-Nbs except O7 bound specifically to T cells upon PHA-L-mediated activation, comparable to the anti-hOX40 mAb. Notably, no binding prior to stimulation was observed (**Figure 4B,C**). These data indicated that O12, O18 and O19 bound endogenous hOX40 exclusively on activated T cells.

**Fig. 4:**
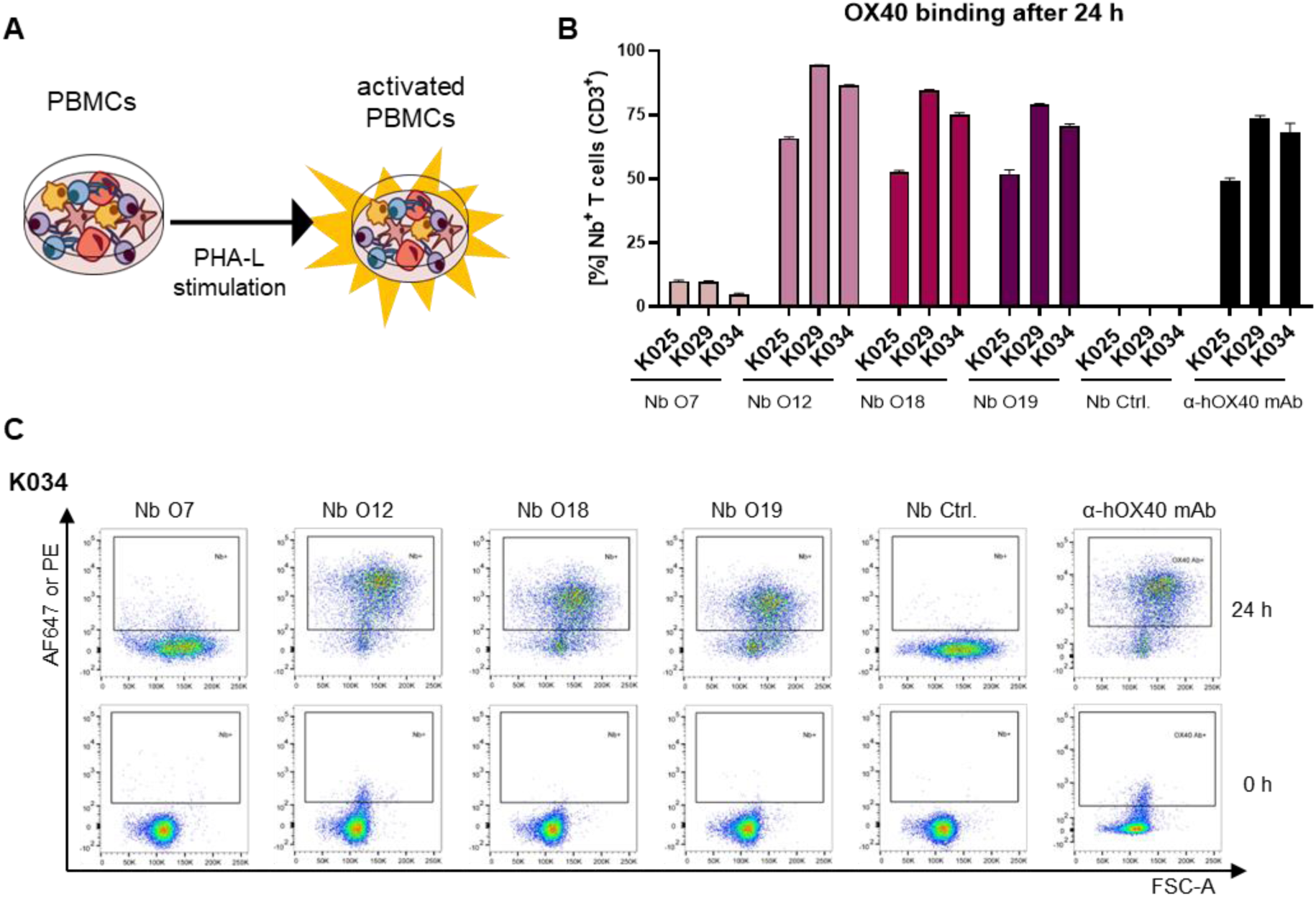
Validation of hOX40-Nb binding to activated T cells. (**A**) Schematic outline of activation of human peripheral blood mononuclear cells (hPBMCs) by phytohaemagglutinin L (PHA-L) stimulation. (**B**) Flow cytometry analysis of hOX40-Nbs staining on CD3^+^ hPBMCs from three different donors (K025, K029 and K034) after 24 h of PHA-L stimulation shown as bar graph. Data are presented as mean ± SD of three technical replicates. (**C**) Exemplary results of flow cytometry analysis of CD3^+^ hPBMCs derived from donor K034 stained with AF647-labeled hOX40-Nbs, an unspecific Nb (Nb. Ctrl.) or an PE-labeled anti-hOX40 mAb before (0 h, lower panel) and after (24 h, upper panel) PHA-L stimulation.

### Agonistic and antagonistic effects of Nbs on OX40 signaling

Targeting OX40, can trigger strong immune responses (41). Therefore, we next investigated whether binding of the Nbs exerts agonistic or antagonistic effects on OX40-mediated signaling by using a genetically engineered Jurkat T cell-based bioassay. These effector cells express hOX40 and contain a luciferase reporter driven by a response element downstream of the OX40 signaling axis. Non-stimulated OX40 effector cells exhibited a weak luminescent signal, which was not further enhanced by addition of increasing concentrations of Nbs O12, O18, and O19. However, when using OX40L as positive control, we observed a strong concentration-dependent induction of NF-kB promotor activity, reflected by an increasing luminescence signal, as expected (**Figure 5A**). Next, we used this assay to analyze a possible competition between OX40L and Nbs for hOX40 binding. Therefore, we preincubated the OX40 effector cells with serial dilutions of Nbs ranging from 0.13 µM to 0.002 nM before adding OX40L at the saturation concentration of 0.12 µM. In this setting, we observed a reduction in luminescence in the presence of Nb O12 and, to a minor extent, of O18, whereas pre-incubation with O19 did not have any effect on OX40L-mediated induction of OX40 signaling (**Figure 5B**). These results were consistent with a BLI-based competition assay in which we monitored the binding of OX40L to recombinant hOX40 in the presence or absence of Nbs (**Supplementary Figure S6**). From these findings, we concluded that none of the Nbs augment OX40-mediated signaling. However, O12 and, to a minor degree, O18, which both target domain 1 of hOX40, competed with the binding of the natural ligand OX40L and could therefore potentially exert an antagonistic effect.

**Fig. 5:**
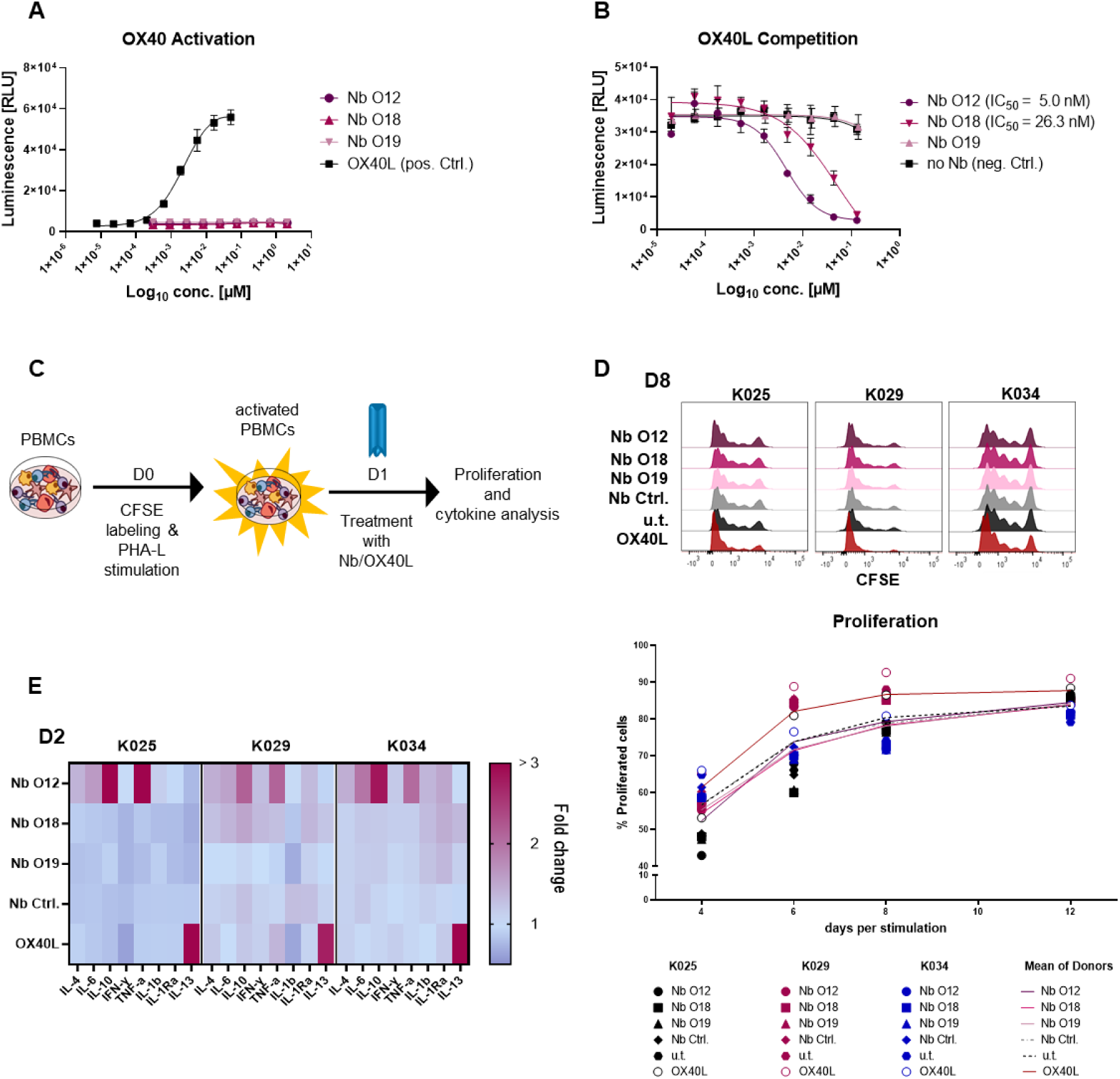
Analysis of hOX40-Nbs on OX40 signaling. (**A, B**) Assessment of agonistic or antagonistic activities of hOX40-Nbs on OX40 signaling in a cell-based OX40 bioassay. (**A**) For determining agonistic effects OX40 effector cells were treated for 5 h with a serial titration of Nbs O12, O18, O19 or OX40L as positive control (pos. Ctrl.) followed by luminescence detection using Bio-Glo reagent. Data are shown as a three-parameter logistic regression dose-response curve based on the mean ± SD of three technical replicates (n = 3). (**B**) For analysis a potential OX40L competition, OX40 effector cells were preincubated with serial dilutions of Nbs ranging from 0.13 µM to 0.002 nM before adding OX40L at the saturation concentration of 0.12 µM followed by luminescence detection using Bio-Glo reagent. Three-parameter logistic regression dose-response curves based on the mean ± SD of three technical replicates showed an antagonistic effect of Nb O12 and O18 with IC50 values of ∼ 5.0 nM or 26.3 nM, respectively. (**C**) Schematic workflow for testing the impact of Nbs on T cell proliferation and cytokine release. hPBMCs of three donors (K025, K029, K034) were CFSE-labeled and stimulated with 5 µg/mL PHA-L. After 24 h, hPBMCs were treated with 0.5 µM hOX40-Nbs, unspecific Nb (Nb Ctrl.), OX40L or left untreated (u.t.). Proliferation at days 4, 6, 8 and 12 and cytokine release at day 2, 4 and 8 were monitored. (**D**) Proliferation was analyzed by flow cytometry (CFSE-low/negative fraction), exemplary shown for day 8 (D8, upper panel). Mean percentages of all three donors are shown as plain or dotted lines (lower panel). (**E**) Determination of cytokines secreted after treatment with hOX40-Nbs displayed as a heat map, exemplary shown for day 2 (D2) after treatment. Values are shown as fold change compared to the untreated control based on the mean of three technical replicates.

### Impact of hOX40-Nbs on proliferation and cytokine release of immune cells

To further explore possible effects of hOX40-Nb binding on T cells, we next investigated its influence on proliferation and cytokine release in hPBMCs. Therefore, hPBMCs from three donors (K025, K029 and K034) were labeled with carboxyfluorescein succinimidyl ester (CFSE) followed by induction of hOX40 expression by PHA-L stimulation for 24 h. After confirming a successful activation and hOX40 expression by flow cytometry (**Supplementary Figure S7A,B**), hPBMCs were either left untreated or incubated with Nbs O12, O18, and O19 or PEP-Nb (42) (Nb Ctrl.) as a negative control (each 0.5 µM). For targeted hOX40 stimulation 0.5 µM OX40L was used (**Figure 5C**). Cell proliferation was monitored on days 4, 6, 8 and 12 by flow cytometry (**Figure 5D**). The results revealed similar proliferation profiles in the hPBMC samples from the same donor upon Nb treatment compared to the untreated samples, while treatment with OX40L induced a ∼10% increase at day 6 and 8 in proliferation (**Figure 5D; Supplementary Figure S7C,D**). In addition, we investigated effects of hOX40-Nbs on the release of cytokines. Therefore, we determined the concentration of a panel of pro- and anti-inflammatory cytokines (**Supplementary Table S2**) in the supernatant of samples collected on days 2, 4, and 8 using a previously reported microsphere-based sandwich immunoassay (11). While Nb O18 and O19 showed only minor effects on cytokine release compared to the untreated control or samples incubated with a non-specific Nb, a significant increase of the NFκ-driven cytokines TNF and IL-6 and of the Th2 cytokines IL-4, and IL-10 upon treatment with O12 was observed. Interestingly, the cytokine release of O12-treated samples differed from samples treated with OX40L, in which only elevated levels of IL-13 were observed (**Figure 5E, Supplementary Figure S8**).

### hOX40-Nb for in vivo imaging

For *in vivo* OI, we chose the fluorescently-labeled Nb O18 (O18_AF647_) as it showed the strongest binding to cellular exposed hOX40 and did not modulate T cell function. CD1 nude mice with subcutaneous HT1080-hOX40 or HT1080-WT tumors were intravenously (*i.v.*) injected with 5 μg of O18_AF647_ and non-invasively *in vivo* investigated by OI over 6 h. The signal intensity (SI) of O18_AF647_ in both the HT1080-hOX40 and the HT1080-WT tumors peaked within 5 min after injection. While the SI continuously decreased over time in the HT1080-WT tumors, it remained stable in the HT1080-hOX40 tumors between 3 h and 6 h post injection, indicating target-specific accumulation after the initial clearing phase of O18_AF647_. Importantly, we determined an increased O18_AF647_-related SI in the HT1080-hOX40 tumors compared to HT1080-WT tumors at all imaging time points with the greatest difference 6 h post injections of ∼7-fold (**Figure 6A, B**). Finally, mice were sacrificed, and the presence of O18_AF647_ within the explanted tumors was analyzed by *ex vivo* OI. Consistent with the *in vivo* data, HT1080-hOX40 tumors exhibited a significantly higher uptake (SI) when compared to HT1080-WT control tumors, indicating a specific binding of the Nb O18 to its target antigen and a favorable signal-to-background ratio for this Nb-derived immunoprobe (**Figure 6C**). These findings demonstrated a specific binding for O18_AF647_ *in vivo*, highlighting its potential as a promising tool for noninvasive monitoring of OX40 expression in immune diagnostics.

**Fig. 6:**
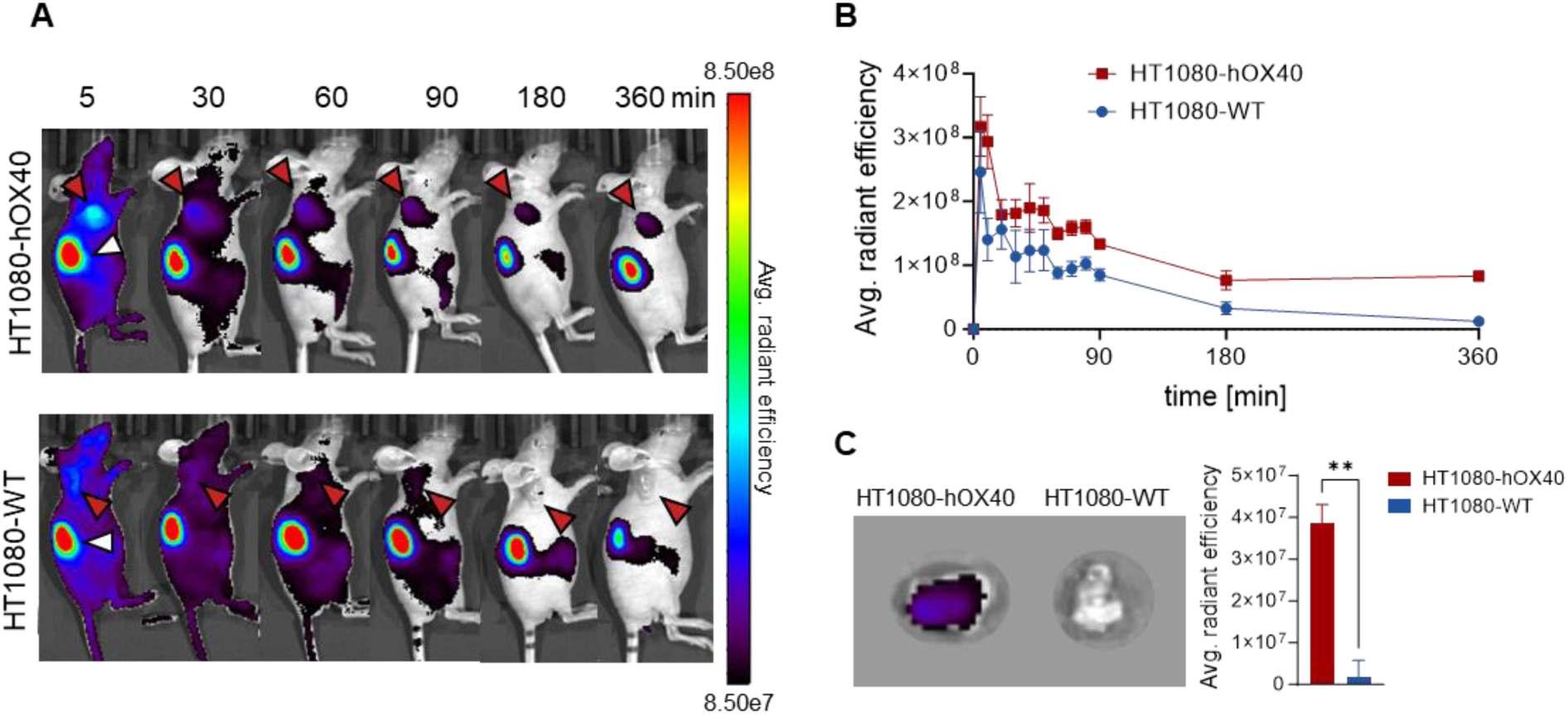
***In vivo* optical imaging with Nb O18AF647** *In vivo* optical imaging (OI) with O18AF647 in HT1080-hOX40 and HT1080-WT tumor bearing mice. 5 µg of O18AF647 were administered intravenously (i.v.) to CD1 nude mice subcutaneously injected with human HT1080-hOX40 or HT1080-WT at the right upper flank. Tumor biodistribution was monitored by repetitive OI measurements over the course of 6 h (**A**) Acquired images of different measurement time points of one representative O18AF647 injected mouse with a HT1080-hOX40 (top) or HT1080-WT (bottom, control). Red arrows indicate the tumor localization at the right upper flank. The kidney is marked with a white arrow at the 5 min time point. (**B**) Quantification of the fluorescence signal from the tumors (n = 3 per group, arithmetic mean of the average radiant efficiency ± SEM) determined at indicated time points. (**C**) Representative *ex vivo* OI of harvested tumor (left) and organ quantification of O18AF647 in HT1080-hOX40 and HT1080-WT tumors (n = 3 per group, arithmetic mean ± SEM; unpaired t test, p = 0.0031).

## Discussion

hOX40 is a recognized theranostic marker that is relevant for both diagnostic and therapeutic applications in the emerging field of immunotherapies (43). On tumor infiltrating T cells, expression of hOX40 correlates with a beneficial outcome and overall survival of patients with solid tumors such as colorectal cancer, cutaneous melanoma, non-small cell lung cancer and ovarian cancer (44–48). In the context of IMIDs such as rheumatoid arthritis, systemic lupus erythematosus or ulcerative colitis, increased expression of hOX40 often correlates with disease activity and severity (49, 50). Building on this potential, anti-OX40-specific mAbs were developed to monitor OX40 expression on activated T cells in preclinical mouse models and used in noninvasive medical imaging of OX40 in cancer vaccination (29), T cell response to glioblastoma (30), acute graft-versus-host disease (31), and rheumatoid arthritis (32).

Due to their unique properties, including specific binding, rapid and deep tissue penetration, short systemic half-life and low immunogenicity, Nbs have emerged as promising building blocks for the development of next-generation imaging probes (37) evidenced by an increasing number of preclinical and first in-human clinical trials (51–54). Here, we developed the first hOX40-specific Nbs as novel probes to specifically address activated T cells. As we focused on developing these binders as potential *in vivo* imaging probes, we aimed for Nbs that have minimal to no effects on OX40 signaling. In total, we identified four hOX40-Nbs with high binding affinity and long-term stability. Epitope mapping categorized the Nbs into two groups addressing either domain 1 or domain 3 of hOX40. However, a precise molecular insight into the recognized structural epitopes remains to be investigated. A more detailed analysis, e.g. by hydrogen-deuterium exchange mass spectrometry (HDX-MS), as already described for other Nbs (11, 13, 55), could facilitate the identification of Nbs that can be used for combinatorial binding, e.g., with OX40 antibodies currently in therapeutic development (56, 57). Three of the selected hOX40-Nbs (O12, O18 and O19) specifically bound to physiologically expressed hOX40 on the surface of activated T cells, whereas none of the selected hOX40-Nbs induced OX40 signaling or affected T cell proliferation. Interestingly, although no agonistic effect was observed, *in vitro* studies with O12 resulted in an increased release of NFκB induced cytokines and Th2 cytokines, which may be caused by a cross-reactivity of this Nb with other members of the tumor necrosis factor receptor superfamily (TNFRSF) or induction of an immune response by an unknown mechanism. For *in vivo* imaging applications, we chose Nb O18 as a lead candidate due to its strong affinity for recombinant and cellularly-exposed hOX40. Apparently, site-directed functionalization employing C-terminal sortagging and DBCO-mediated click chemical conjugation to AF647 (11, 58) did not affect its binding properties. OI in a mouse xenograft model showed rapid accumulation of fluorescently labeled O18 to HT1080-hOX40 tumors and sustained binding over a prolonged period of time, indicating high *in vivo* binding functionality with a low off-rate.

In summary, the specific detection of activated T cells can support therapy monitoring and patient selection for personalized immunotherapies such as immune checkpoint inhibitor therapies, where a high number of activated T cells within the TME is beneficial. Furthermore, our approach enables the evaluation of responses to cancer vaccination or cytokine therapy and provides information on the activation status of CAR-T cells. In this context, several probes have been developed to visualize T cell activation markers, including imaging probes based on peptides that bind granzyme B (59), mAbs against ICOS (15), IFN-y (60), murine OX40 (29) or a CD69 targeting affibody as an early phenotypic activation marker (16, 61). Our first hOX40-specific Nbs described here now complete this list. Due to their high binding functionality and unique properties in terms of tissue penetration and improved signal-to-noise ratio without unwanted modulation of OX40 signaling, we anticipate that our hOX40-Nbs will facilitate the visualization of even small numbers of OX40-activated T cells. Because of their fast pharmacokinetics, it is conceivable that these Nbs will enable earlier imaging timepoints and therefore, the usage of shorter-lived isotopes, e.g., ^18^F, thereby reducing patients’ radiation exposure and allowing more longitudinal imaging to assess dynamic changes in the T cell composition within the TME, IME and the lymphatic organs. In perspective, the hOX40-Nbs can not only be used to detect activated T cells in cancer lesions, but also for the applications in the diagnosis of IMIDs like rheumatoid arthritis (32) or graft-versus-host disease (31).

## Data availability

The data that support the findings of this study are available from the corresponding author upon reasonable request.

## Ethics Statement

With the approval of the Government of Upper Bavaria (approval number: 55.2-1-54-2532.0-80-14). All animal experiments were carried out in accordance with the German Animal Welfare Act and with consent of regulatory authorities (Regierungspräsidium Tübingen).

## Authorship Contributions

D.F., M.K., B.P. and U.R. designed the study. S.N. and A.S. immunized the animal. D.F., T.W. and P.K. performed Nb selection. D.F., T.W., B.T. and P.K. performed biochemical, cell biological characterization and functionalization of Nbs. C.G. provided hPBMC samples and expertise for T cell assays. M.F., M.J., and N.S.M. analyzed the Nb effects on cytokine expression. S.B. and D.S. performed *in vivo* imaging and analysis. M.K., B.P. and U.R. supervised the study. D.F. and U.R. drafted the article. All authors contributed to the article and approved the submitted version.

## Acknowledgements

This work received financial support from the State Ministry of Baden-Wuerttemberg for Economic Affairs, Labour and Tourism (Grant: Predictive diagnostics of immune-associated diseases for personalized medicine. FKZ: 35-4223.10/8). This work was further supported by the Deutsche Forschungsgemeinschaft (DFG, German Research Foundation, Germanýs Excellence Strategy-EXC2180-390900677) and the Werner Siemens-Foundation.

## Materials & Methods

### Expression constructs

hOX40-encoding DNA (GenBank accession: NM_003327.3) was synthesized and cloned into NheI and EcoRI site of pcDNA3.1(+) by GenScript Biotech. The vector backbone was changed by cutting with the restriction enzymes EcoRI and BstBI into a backbone comprising an internal ribosomal entry site (IRES) and genes for eGFP as reporter and Blasticidin S deaminase for antibiotics resistance from the expression construct as described previously (11). For the generation of hOX40 domain deletion mutant expression constructs hOX40ΔD1 (aa 66-277), hOX40ΔD1-2 (aa 108-277), hOX40ΔD1-3 (aa 127-277) of UniProtKB P43489, respective fragments were amplified (**Supplementary Table S3**) and genetically fused N-terminally to a SPOT-Tag (62). DNA encoding for murine OX40 (mOX40) was purchased from Sino Biological (Catalog Number MG50808-NM).

### Stable cell line generation and culturing

U2OS cells (ATCC) were cultured according to standard protocols in Dulbecco’s modified Eagle’s medium (DMEM), supplemented with 10 % (v/v) FCS and 1 % (v/v) penicillin/streptomycin (all Thermo Fisher Scientific). Cultivation conditions were 37°C and 5 % CO_2_ atmosphere in a humidified incubator and passaged using 0.05 % trypsin-EDTA (Thermo Fisher Scientific). Transfection of plasmid DNA was performed using Lipofectamine 2000 (Thermo Fisher Scientific) according to the manufacturer’s protocol. To generate U2OS cells stably overexpressing hOX40 on their surface (U2OS-hOX40), 24 h after transfection, selection pressure by 5 µg/mL Blasticidine S (Sigma Aldrich) was applied for a period of two weeks. After single cell separation, monoclonal cells were analyzed for hOX40 expression using live-cell fluorescence microscopy.

### Animal immunization and hOX40-Nb library generation

The alpaca immunization was performed with the approval of the Government of Upper Bavaria (approval number: 55.2-1-54-2532.0-80-14). Two alpacas (*Vicugna pacos*) were immunized using the extracellular part of recombinant hOX40 (hOX40 AA Leu 29 – Ala 216) produced in human HEK293 cells (Acrobiosystems). During a period of 91 days, the animals were vaccinated six times at day 0, 21, 28, 35, 49, 87. The initial vaccination was performed with 560 µg followed by five booster injections each consisting of 280 µg hOX40 with Adjuvant F (Gebru). After the 91-day period, lymphocytes were isolated from ∼200 mL of blood performing Ficoll gradient centrifugation with lymphocyte separation medium (Carl Roth) and total RNA was extracted by NucleoSpin® RNA II (Macherey&Nagel). The mRNA was subsequently transcribed into cDNA by the First Strand cDNA Synthesis Kit (GE Healthcare). The Nb repertoire was isolated as described in three subsequent PCR reactions using the following primer combinations: (1) CALL001 and CALL002, (2) forward primers FR1-1, FR1-2, FR1-3, FR1-4, and reverse primer CALL002, and (3) forward primers FR1-ext1 and FR1-ext2 and reverse primers FR4-1, FR4-2, FR4-3, FR4-4, FR4-5, and FR4-6 introducing SfiI and NotI restriction sites (11) (**Supplementary Table S3**). This enables subcloning of Nb library into the pHEN4 phagemid vector (63)

### Nb screening

The selection of hOX40-specific Nbs was performed by two consecutive rounds of phage display against immobilized recombinant antigen. For this purpose, electrocompetent TG1 *E. coli* bacteria were transformed with the hOX40-Nb library in pHEN4 and infected with M13K07 helper phages leading to the generation of hOX40-Nb presenting phages. 1 x 10^11^ phages were enriched by adsorption to streptavidin or neutravidin plates (Thermo Fisher Scientific) coated with hOX40 (5 µg/mL), biotinylated by Sulfo-NHS-LC-LC-Biotin (Thermo Fisher Scientific) in 5 molar excess at ambient temperature for 30 min and purified using Zeba^TM^ Spin Desalting Columns 7 K MWCo 0.5 mL (Thermo Fisher Scientific) according to manufacturer’s protocol. Antigen and phage blocking was performed with 5% milk in PBS-T during the first round or BSA in the second. Washing stringency was increased with each panning round and elution of bound phages was performed by 100 mM triethylamine pH 10 (TEA, Roth), followed by neutralization with 1 M Tris/HCl pH 7.4. For phage rescue, TG1 bacteria were infected with the eluted phages during their exponential growth phase, spread on selection plates for subsequent selection rounds and incubated at 37°C overnight. Enrichment of antigen-specific phages was monitored by counting colony forming units (CFUs).

### Whole-cell phage ELISA

Monoclonal phage ELISA was executed in a whole cell setting. Individual clones were picked, and phage production was induced as described above. For antigen presentation U2OS-hOX40 cells or wild type (wt) U2OS for background determination were seeded in a density of 2 × 10^4^ cells per well in 100 µL in 96-well cell culture plates (Corning) coated with poly-L-lysine (Sigma Aldrich) and grown overnight. The next day, 70 µL of phage supernatant was added to each cell type and incubated at 4°C for 3 h. Cells were washed 5 × with 5% FCS in PBS, followed by incubation with M13-HRP-labeled detection antibody (Progen, 1:2000 Dilution) for 1 h and washed again 3 × with 5% FCS in PBS. For the final detection, Onestep ultra TMB 32048 ELISA substrate (Thermo Fisher Scientific) was added to each well and incubated until the color changed. The reaction was stopped with 100 µL of 1 M H_2_SO_4_ and the signal was detected with the Pherastar plate reader at 450 nm. Phage ELISA-positive clones were defined by a 2-fold signal above U2OS-WT control cells.

### Protein expression and purification

For production, hOX40-Nbs were cloned into pHEN6 vector (63), expressed in XL-1 and purified using immobilized metal affinity chromatography (IMAC) and size exclusion chromatography according to standard procedures as previously described (11). Sortase A pentamutant (eSrtA) in pET29 was a gift from David Liu (Addgene plasmid # 75144) and expressed and purified as published (64). The quality of all purified proteins was analyzed via standard SDS-PAGE under denaturizing and reducing conditions (5 min, 95°C in 2x SDS-sample buffer containing 100 mM Tris/HCl, pH 6.8; 2 % (w/v) SDS; 5 % (v/v) 2- mercaptoethanol, 10 % (v/v) glycerol, 0.02 % bromphenole blue). Proteins were visualized by InstantBlue Coomassie (Expedeon) staining or alternatively by immunoblotting transferring proteins to nitrocellulose membrane (GE Healthcare, Chicago, IL, USA) and detection using a primary anti-Penta-His antibody (Qiagen) and secondary donkey anti-mouse AF647 antibody (Invitrogen) on a Typhoon Trio scanner (GE-Healthcare, excitation 633 nm, emission filter settings 670 nm BP30).

### Biolayer interferometry (BLI)

The binding kinetics analysis of hOX40-Nbs was performed using the Octet RED96e system (Sartorius) applying manufacturer’s recommendations. Therefore, 5 µg/mL of biotinylated hOX40-Nbs diluted in Octet buffer (PBS, 0.1 % BSA, 0.02 % Tween20) were immobilized on streptavidin coated biosensor tips (SA, Sartorius) for 30 s and unbound Nb was washed away. For the association step, a dilution series of hOX40 ranging from 0.2 nM – 320 nM were applied for 300 s followed by dissociation in Octet buffer for 720 s. Each concentration was normalized to a reference applying Octet buffer only for association. Data were analyzed using the Octet Data Analysis HT 12.0 software applying the 1:1 ligand-binding model and global fitting. For epitope binning 5 µg/mL of each Nb, except O7 due to its inappropriate dissociation behavior, was immobilized to SA tips and the association was performed with a premixture of OX40 (100 nM) and an excess of unbiotinylated second Nb (1000 nM). By analyzing the binding behavior of the premixture, conclusions about shared epitopes were drawn. To determine a potential competition of Nbs with the natural OX40 ligand OX40L, OX40L was biotinylated and 10 µg/mL were immobilized to the SA tips. Premixture of hOX40 with a ten-time molar excess of each Nb was applied for the association step.

### Live-cell immunofluorescence

U2OS-hOX40 cells, U2OS-WT or U2OS cells transiently expressing hOX40 domain deletion mutants or murine OX40 were plated at a density of 1 x 10^4^ cells per well in 100 µL of a µClear 96-well plate (Greiner Bio One, cat. #655090) and cultivated overnight at standard conditions. The next day, cells were stained with 2 µg/mL Hoechst33258 (Sigma Aldrich) for nuclear staining in live-cell visualization medium DMEMgfp-2 (Evrogen, cat. #MC102) supplemented with 10 % FCS for 30 min at 37°C. Afterwards 10 -1000 nM fluorescently labeled hOX40-Nbs, an unspecific Nb (Mock) or an OX40 antibody (positive control) were added and incubated for 30 min at 4°C. Staining solution was replaced by live-cell visualization medium DMEMgfp-2 with 10 % FCS and images were acquired with a MetaXpress Micro XL system (Molecular Devices) at 20 x magnification.

### Stability analysis

To assess the thermal stability of the Nbs, nanoscale differential scanning fluorimetry (nanoDSF) with the Prometheus NT.48 device (Nanotemper) was performed. Freshly thawed hOX40-Nbs were diluted to 0.25 mg/mL in PBS and measured at time point d0 and after an incubation period of ten days at 37°C (d10) using standard capillaries. A thermal gradient ramping from 20°C to 95°C was applied while measuring fluorescence ratios (F350/F330) and light scattering. Using PR. ThermControl v2.0.4 the melting (T_M_) and aggregation (T_Agg_) temperatures were determined.

### Fluorescent labeling of Nanobodies

For sortase A based coupling of 50 μM Nb were added to 250 μM sortase peptide (H-Gly-Gly-Gly-propyl-azide synthesized by Intavis AG) and 10 μM sortase A both dissolved in sortase buffer (50 mM Tris, and 150 mM NaCl, pH 7.4 at 4°C) and reaction was started by adding 10 mM CaCl_2_ for 4 h at 4°C. To avoid reverse sortase reaction, sortase A and uncoupled Nb were removed by Ni-NTA affinity chromatography. The coupled Nbs were concentrated and residual peptide was depleted using Amicon ultra-centrifugal filters MWCO 3 kDa. Taking advantage of SPAAC (strain-promoted azide-alkyne cycloaddition) click chemistry reaction fluorescent labeling was performed by incubating azide-coupled Nbs with 2-fold molar excess of DBCO-AF647 (Jena Bioscience) for 2 h at room temperature. Subsequent dialysis (GeBAflex-tube, 6-8 kDa, Scienova) led to removal of excess of DBCO-AF647. As final polishing step a hydrophobic interaction chromatography (HIC, HiTrap Butyl-S FF, Cytiva) was performed to deplete unlabeled Nb. The final products were analyzed via SDS-PAGE and spectrophotometry.

### OX40 bioassay

The OX40 bioassay kit (Promega) was used to determine a potential agonistic activity of Nbs O12, O18 and O19 according to manufactureŕs instructions. On the day before assay, thaw-and-use OX40 effector cells (Promega) were thawed and seeded into the inner 60 wells of two white 96 well assay plates cultured in assay buffer (RPMI1640 with 5% FBS) at standard conditions overnight. The next day OX40L and Nbs were serially diluted (OX40L: 50 – 0.0008 nM; Nbs: 2000-0.3 nM) in assay buffer, and 20 μL of the diluted recombinant proteins were added to the assay plate in triplicates. The assay plate was incubated at 37°C, 5 % CO_2_ for 5 h. Afterwards, the assay plate was equilibrated to ambient temperature for 10 min. For detection of agonistic function, 75 µL of Bio-GloReagent was added to all wells. The assay plate was incubated at room temperature for 5 min, and luminescence was measured using a Tecan M2000 plate reader. The average relative luminescence unit (RLU) was calculated for each dilution. The average RLU data were plotted against the different concentrations of OX40L and hOX40-Nbs Nbs. To test antagonistic properties of the Nbs, the OX40 bioassay was transformed into a competition assay. For this purpose, the cells were pre-incubated with a serial dilution of hOX40-Nbs (0.13 µM to 0.002 nM) for one hour, followed by a 5 h incubation period with 0.12 µM OX40L.

### Peripheral blood mononuclear cells isolation and start of culture

Human peripheral blood mononuclear cells (hPBMCs) were isolated as described in (11). In brief, fresh mononuclear blood cell concentrates were obtained from healthy volunteers at the ZKT Tübingen gGmbH. Participants gave informed written consent and the studies were approved by the ethical review committee of the University of Tübingen, projects 156/2012BO1 and 713/2018BO2. Blood products were diluted with PBS 1x (homemade from 10x stock solution, Lonza, Switzerland) and PBMCs were isolated by density gradient centrifugation with Biocoll separation solution (Biochrom, Germany). PBMCs were washed twice with PBS 1x, counted with a NC-250 cell counter (Chemometec, Denmark), resuspended in heat-inactivated (h.i.) fetal bovine serum (FBS) (Capricorn Scientific, Germany) containing 10% DMSO (Merck) and frozen in aliquots using a freezing container before transfer to nitrogen for long term storage. For the experiments, cells were thawed in Iscovés Modified Dulbeccós Medium (IMDM + L-Glutamin + 25 mM HEPES; Thermo Fisher Scientific) supplemented with 2.5 % h.i. FBS (Thermo Fisher Scientific), 1 % P/S (Sigma-Aldrich), and 50 μM β-mercaptoethanol (β-ME; Merck), washed once, counted, and rested for 1 h at 37°C 5 % CO_2_ in T cell medium (TCM, IMDM + 2 % h.i. FBS + 1x P/S + 50 µM β-ME) supplemented with 1 µg/mL DNase I (Sigma-Aldrich). After resting, cells were washed once again, counted and used for subsequent analysis.

### Validation of hOX40-Nb binding to activated T cells

For the validation of Nb binding to activated T cells, hPBMCs were stained before and after stimulation for 24 h with 5 µg/mL PHA-L in TCM at 37°C 5 % CO_2_. For flow cytometry analysis 2x10^5^ cells per staining condition in FACS buffer (PBS containing 0.02 % sodium azide, 2 mM EDTA, 2% h.i. FBS) were used. Extracellular staining was performed with AF647-labeled hOX40-Nbs or unspecific (Nb Ctrl.) Nb (200 nM), phycoerythrin (PE)-labeled anti-hOX40 mAb (Ber-Act35, BioLegend), CD3 Ab APC-Cy7 (HIT3a, BioLegend), dead cell marker Zombie Violet (BioLegend) and isotype control Abs (BioLegend) each in pretested optimal concentrations by incubation for 30 min at 4°C. Cells were washed twice with FACS buffer and acquired on the same day using a LSRFortessa^TM^ flow cytometer (Becton Dickinson) equipped with the DIVA Software (Becton Dickinson). Final data analysis was performed using the FlowJo10® software (Becton Dickinson).

### T cell proliferation assay

The proliferation behavior of T cells was assessed using a carboxyfluorescein succinimidyl ester (CFSE) based approach. Up to 1×10^8^ cells were labeled with 2.5 µM CFSE (BioLegend) in 1 ml PBS for 20 min according to the manufacturer’s protocol. The cells were washed twice in medium containing 10% h.i. FBS to stop CFSE labeling and stimulated for 24 h with 5 µg/mL PHA-L in TCM at 37°C 5 % CO_2_ in a 48-well cell culture plate with 1.6–2.5×10^6^ cells/well. Induced OX40 expression was validated via flow cytometry. Subsequent to the stimulation, hPBMCs were treated with 0.5 µM of OX40 specific Nbs O12, O18 and O19, a control Nb, OX40L or left untreated and cultured at 37°C and 5 % CO_2_. Concentrations were chosen in a large excess than the expected concentration during clinical application. On days 3, 5 and 7, 2 ng/mL recombinant human IL-2 (R&D, USA) were added. One-third of the culture on day 4, one half of the culture on days 6 and 8, and the remaining cells on day 12 were harvested and counted. Cells from each condition were washed twice with FACS buffer (PBS containing 0.02% sodium azide, 2 mM EDTA, 5% h.i. FBS). Extracellular staining was performed with CD3 Ab APC-Cy7 (HIT3a, BioLegend), dead cell marker Zombie Violet (BioLegend) and isotype control Abs (BioLegend) each in pretested optimal concentrations by incubation for 30 min at 4°C. Cells were washed two times with FACS buffer and acquired on the same day using a LSRFortessaTM flow cytometer (Becton Dickinson) equipped with the DIVA Software (Becton Dickinson). Final data analysis was performed using the FlowJo10® software (Becton Dickinson). The percentage of proliferating T cells was determined by assessment of CFSE negative cells.

### Cytokine release assay

For cytokine release analysis, a set of in-house developed Luminex-based sandwich immunoassays was used. Supernatants from days 2, 4 and 8 of the proliferation assay were frozen at -80°C until cytokine measurements. Levels of IL-1b, IL-1Rα, IL-4, IL-6, IL-8, IL-10, IL-12p70, IL-13, granulocyte-macrophage colony-stimulating factor (GM-CSF), IFN-γ, macrophage chemotactic protein (MCP)-1, macrophage inflammatory protein (MIP)-1b, TNFα, and vascular endothelial growth factor (VEGF) were determined using a those immunoassays each consisting of commercially available capture and detection antibodies and calibrator proteins. All assays were thoroughly validated ahead of the study with respect to accuracy, precision, parallelism, robustness, specificity, and sensitivity (65, 66). Samples were diluted at least 1:4 or higher. After incubation of the prediluted samples or calibrator protein with the capture coated microspheres, beads were washed and incubated with biotinylated detection antibodies. Streptavidin-phycoerythrin was added after an additional washing step for visualization. For control purposes, calibrators and quality control samples were included on each microtiter plate. All measurements were performed on a Luminex FlexMap® 3D analyzer system using Luminex xPONENT® 4.2 software (Luminex, USA). For data analysis, MasterPlex QT, version 5.0, was employed. Standard curve and quality control samples were evaluated according to internal criteria adapted to the Westgard Rules (67) to ensure proper assay performance.

### hOX40-Nbs for in vivo optical imaging

For optical *in vivo* imaging, we labeled the Nb O18 with the fluorophore AlexaFluor647 (O18_AF647_) by sortase-mediated attachment of an azide group followed by click-chemistry addition of DBCO-AF647. To establish hOX40^+^ expressing tumors, 5 x 10^6^ HT1080 cells stably expressing human OX40 (HT1080-hOX40) or 5 x 10^6^ wild type HT1080 (HT1080-WT) cells serving as negative control were resuspended in 50% Matrigel (BD) and 50% PBS and subcutaneously injected into the right upper flank of 7-week-old CD1 nude mice (Charles River Laboratories). When the tumors reached a size of 50 - 100 mm³, HT1080-hOX40 (n=3) or HT1080-WT (n=3) bearing mice were *i.v.* injected with 5 µg of Nb O18_AF647_ and noninvasively *in vivo* investigated by optical imaging (OI). For the *in vivo* measurements, the mice were anaesthetized with 1.5% isoflurane and the body temperature was kept constant at 37 °C using a heating mat. The mice were imaged over the course of 6 h and were sacrificed after the last imaging time point before tumors were explanted for *ex vivo* OI analysis. A bright field image and an image of the fluorescence signal (excitation 640 nm/ emission 680 nm) were recorded using an IVIS Spectrum OI System (PerkinElmer, Waltham, MA, USA). The fluorescence intensities were quantified by drawing regions of interest around the tumor borders and were expressed as average radiant efficiency (photons/s)/(μW/cm^2^) subtracted by the background fluorescence signal using the Living Image software 4.4 (Perkin Elmer). Statistical analyses were conducted with graph pad prism, Version 10. All mouse experiments were performed according to the German Animal Protection Law and were approved by the local authorities (Regierungspräsidium Tübingen).

## Supplementary Information

### Supplementary Figures

**Fig. S1:**
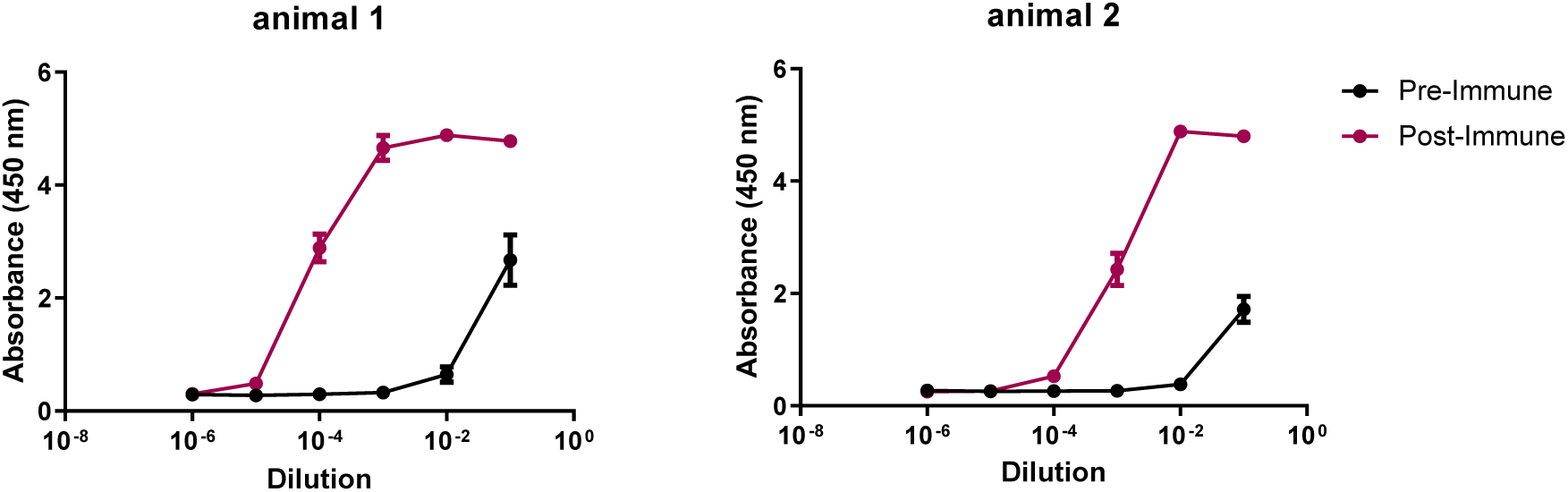
Analysis of seroconversion upon vaccination with hOX40. Analysis of seroconversion upon vaccination with recombinant hOX40. Serum samples of the two vaccinated alpacas (*Vicugna pacos*) were collected before (pre-immune; black data points) and after 63 days of vaccination (post-immune; red data points). To test for the development of hOX40-specific antibodies, a serum ELISA was performed with the indicated dilutions of pre- and post-immune sera in multi-well plates coated with recombinant hOX40. Binding of hOX40-specific antibodies was detected by using an anti-heavy chain antibody conjugated to horseradish peroxidase.

**Fig. S2:**
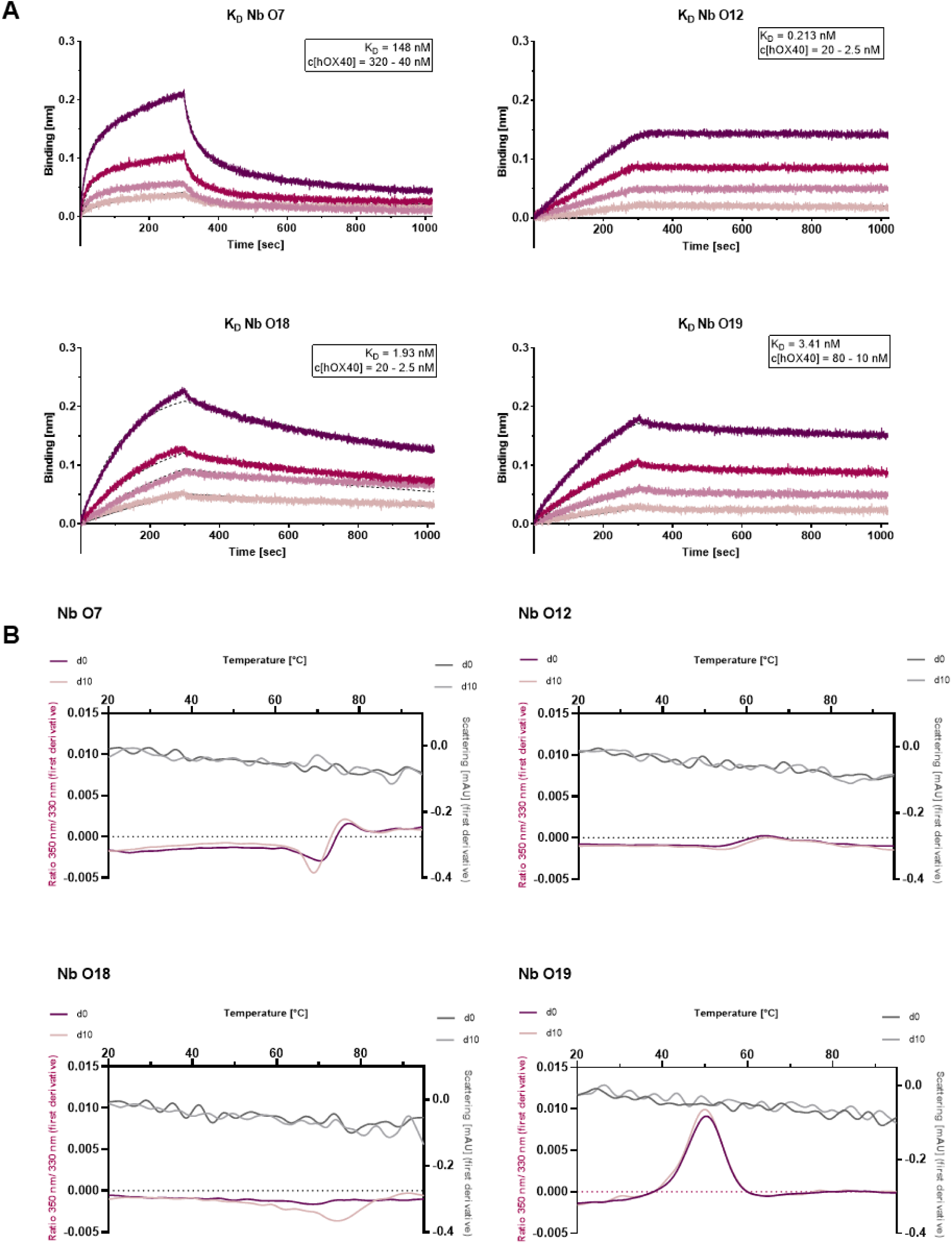
Affinity and stability measurements of hOX40-Nbs. (**A**) Biolayer interferometry (BLI)-based affinity measurements were performed by immobilization of biotinylated hOX40-Nbs on streptavidin sensors. Kinetic measurements were performed using four concentrations (as indicated) of recombinant hOX40 (displayed with gradually lighter shades of color). The indicated binding affinities (KD) were calculated from global 1:1 fits shown as dashed lines. (**B**) Stability analysis of individual hOX40-Nbs using nano scale differential scanning fluorimetry (nanoDSF) displaying fluorescence ratio (350 nm/330 nm) (red) and light scattering (gray) are shown as first derivatives for day 0 (dark shade) and after an accelerated aging period of 10 days at 37°C (light shade).

**Fig. S3:**
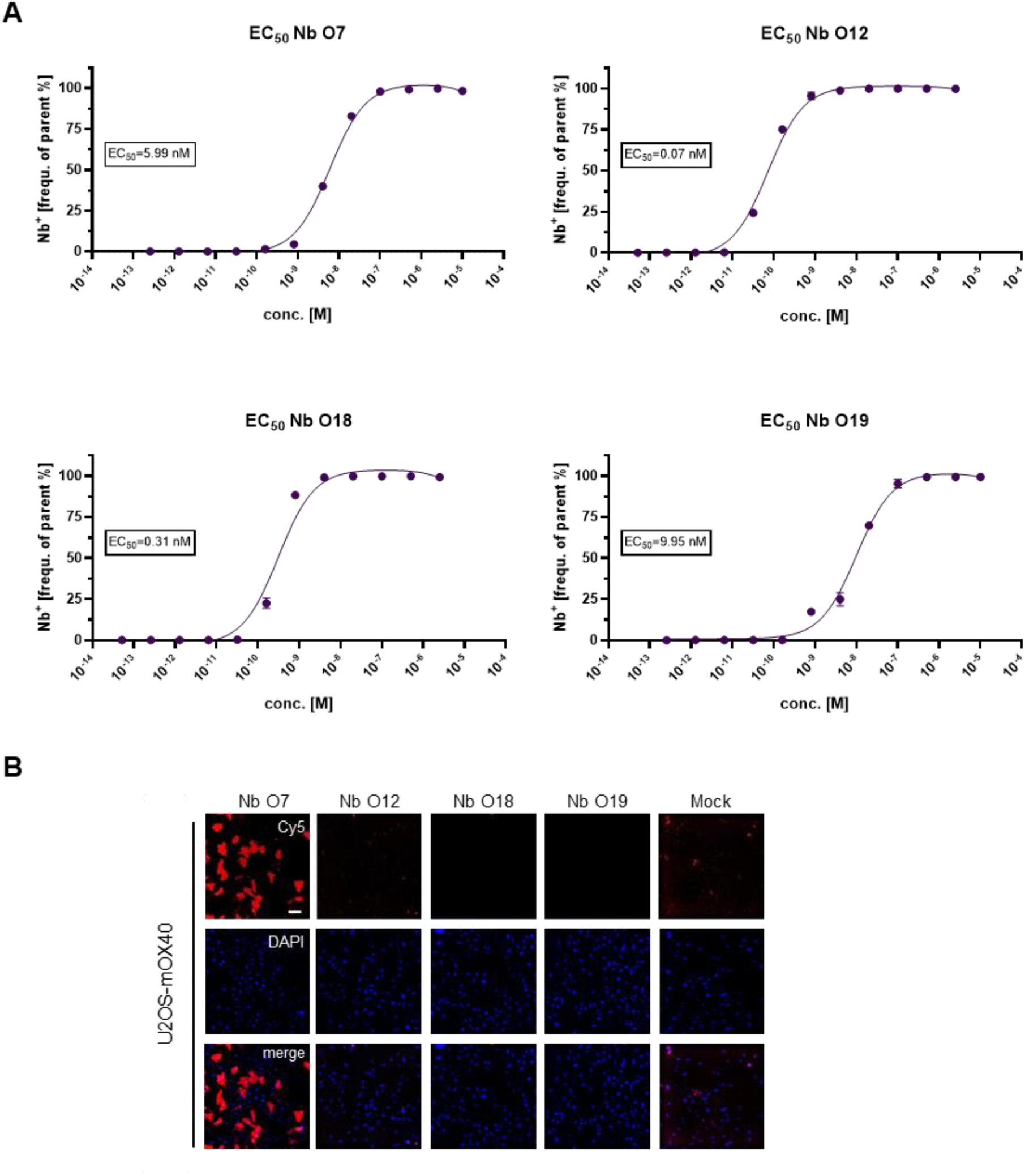
Characterization of cellular binding of hOX40-Nbs. (**A**) Determination of hOX40-Nb binding to cellular expressed hOX40 by flow cytometry. The percentage of positively stained U2OS-hOX40 cells with AlexFluor647-(AF647) labeled hOX40-Nb (frequency of parent) was plotted against indicated concentrations of hOX40-Nbs and indicated EC50 values were calculated from a four-parametric sigmoidal model based on the mean ± SD of three technical replicates (n = 3). (**B**) Cross-reactivity analysis of hOX40-Nbs by immunofluorescence staining. Representative images of U2OS cells transiently expressing murine OX40 (U2OS-mOX40) cells stained with AF647-labeled hOX40-Nbs or an unspecific Nb (Mock) as negative control. Shown are staining with individual Nbs, nuclei staining (Hoechst, blue) and merged signals; scale bar: 50 µm.

**Fig. S4:**
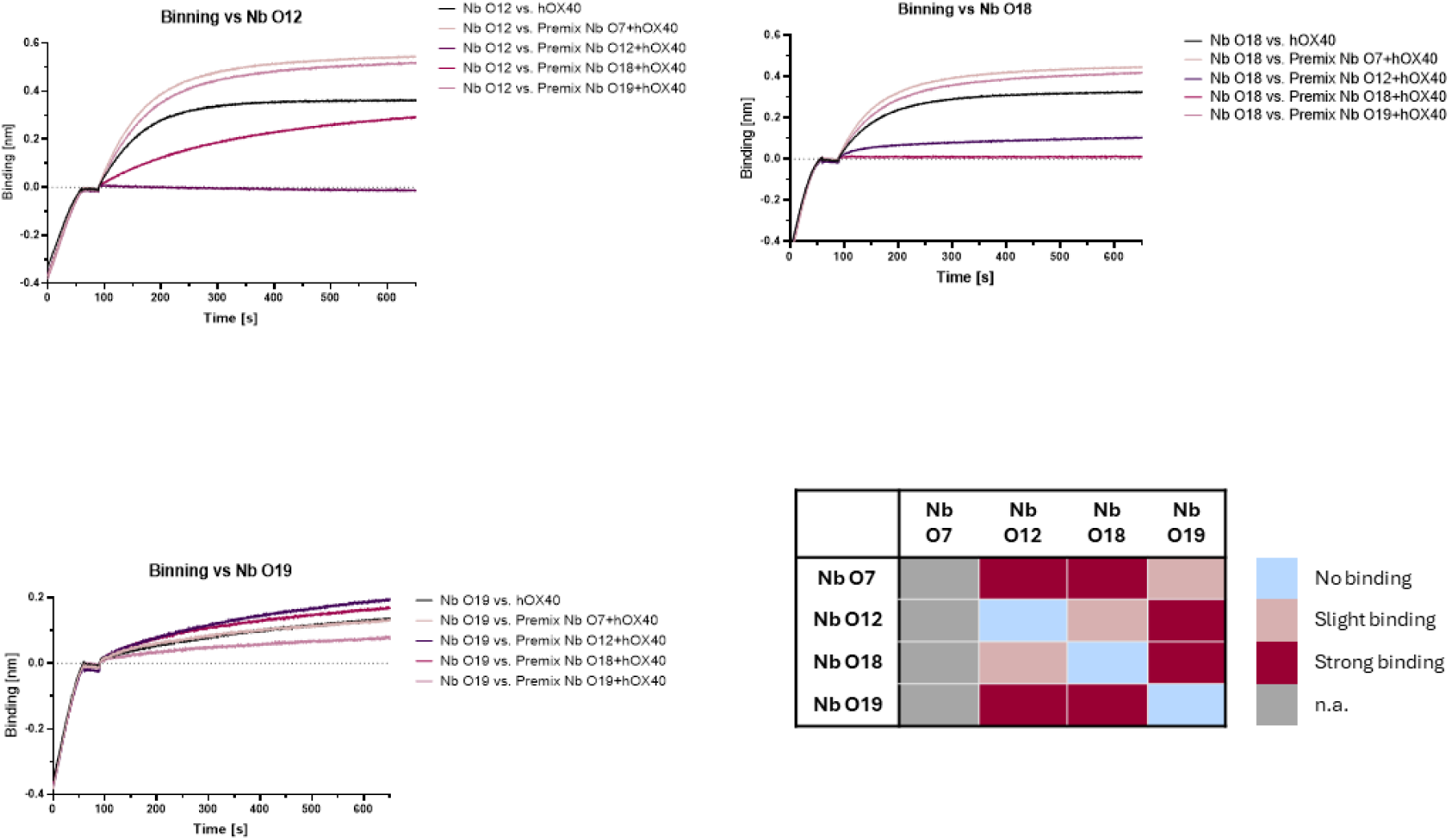
Epitope binning of hOX40-Nbs by biolayer interferometry (BLI) Biotinylated first hOX40-Nb was immobilized on streptavidin biosensors. hOX40 (100 nM) was pre-incubated (premix) in 10-fold excess with designated second hOX40-Nb (as indicated) followed by determining additional association of the hOX40/Nb complex to the immobilized first hOX40-Nb. All sensograms of combinatorial Nb binding to hOX40 on sharing/overlapping epitopes or on different epitopes including a graphical summary of epitope binning analysis are shown.

**Fig. S5:**
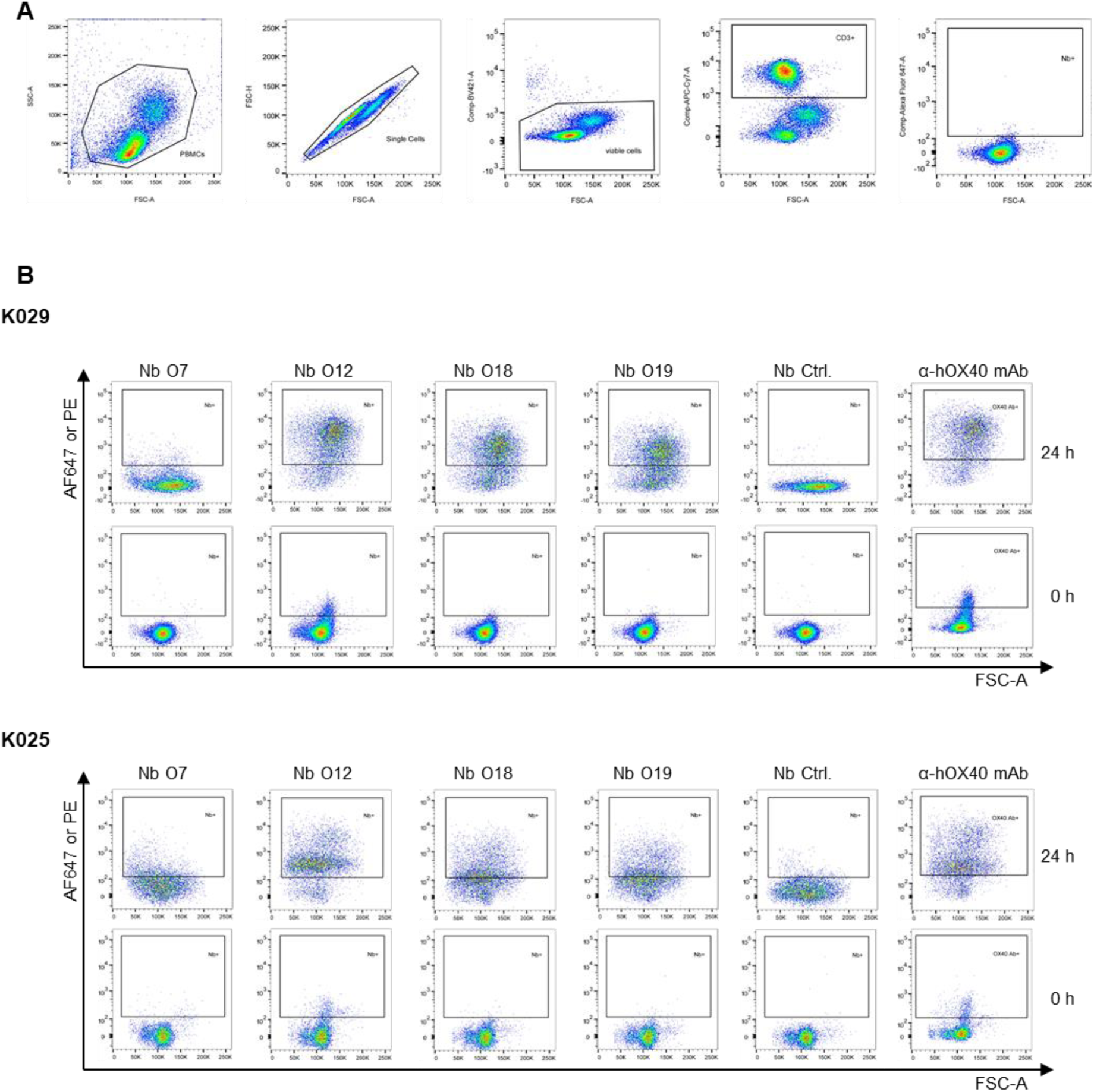
Validation of hOX40-Nb binding to activated T cells. (**A**) Gating strategy for flow cytometry analysis of hOX40-Nb staining of CD3^+^ cells derived from human PBMCs (hPBMC). Isotype control mAbs were used for setting the gates of anti-CD3 mAb and anti-hOX40 mAb (**B**) Flow cytometry analysis of CD3^+^ hPBMCs derived from donors K029 and K025 stained with AF647-labeled hOX40-Nbs, an unspecific Nb (Nb Ctrl.) or phycoerythrin (PE)-labeled anti-hOX40 mAb before (0 h, lower panel) and after (24 h, upper panel) stimulation with 5 µg/mL phytohaemagglutinin L (PHA-L).

**Fig. S6:**
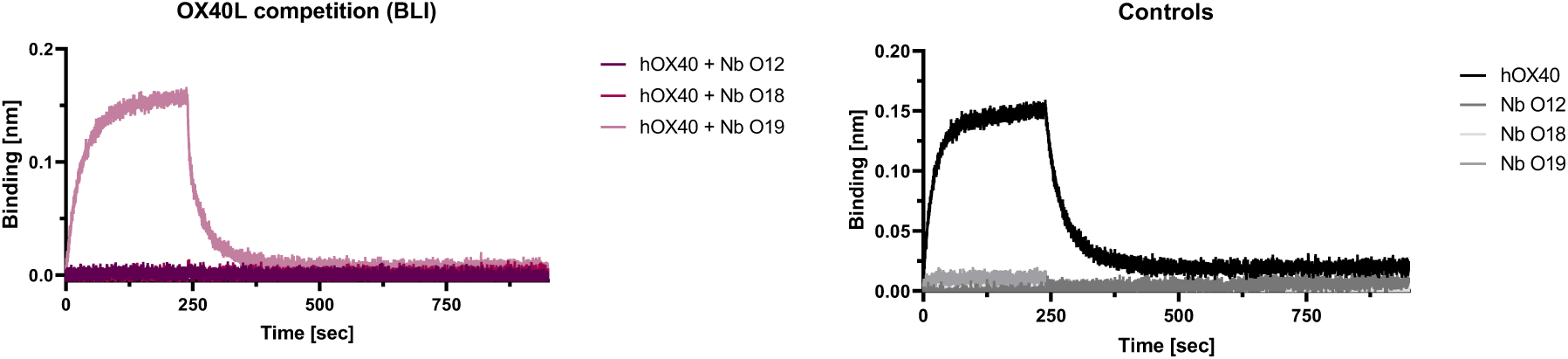
Nb O12 and O18 compete with binding of OX40L to recombinant hOX40. BLI-based OX40L competition assay. OX40L was immobilized to the sensor tips followed by addition of hOX40 either left untreated or preincubated with a 10-fold molar excess of Nb O12, O18 or O19. To exclude artefacts binding of each analyte to hOX40 was also tested alone (Controls).

**Fig. S7:**
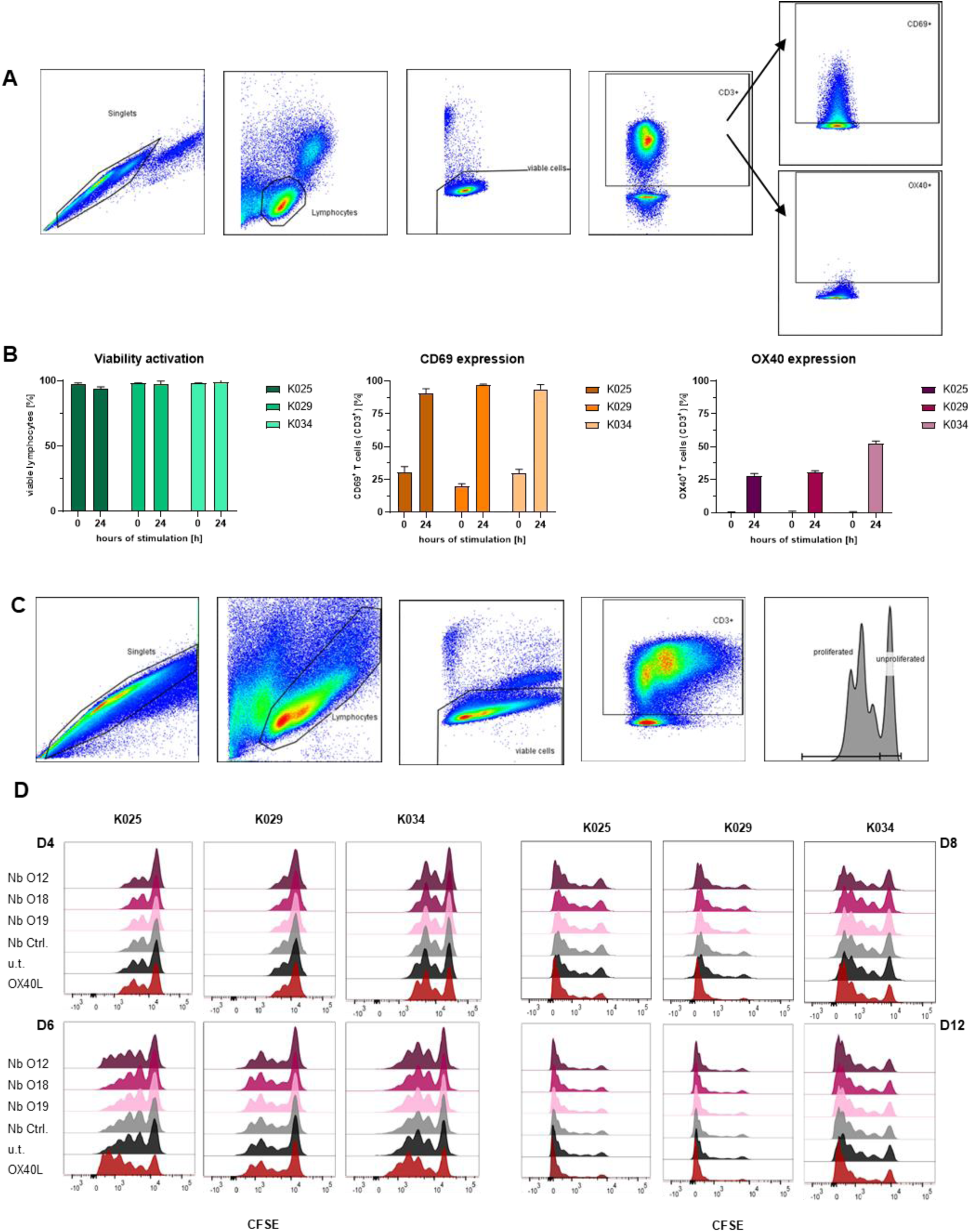
Flow cytometry analysis of effects of hOX40-Nb binding. (**A**) Gating strategy for monitoring successful activation and hOX40 expression on hPBMCs upon PHA-L stimulation (exemplary shown for K034 at day 0). From left to right: single cells, lymphocytes, viable cells, CD3^+^ cells and CD69 expressing (upper gate) or OX40 expressing (lower gate) cells. Isotype control mAbs were used for setting the gates of anti-CD3, anti-hOX40 and anti-CD69 mAbs. (**B**) Viability (left), CD69 expression (mid) and OX40 expression (right) before and after stimulation for the three donors K025, K029 and K034. Data are presented as mean ± SD of three technical replicates (n = 3) (**C**) Gating strategy for T cell proliferation assay (exemplary shown for K034, 4 days after Nb treatment). From left to right: single cells, lymphocytes, viable cells, CD3^+^ cells and CFSE-low/negative cells. Isotype control mAb was used for setting the gates of anti-CD3 mAb. (**D**) Histogram overlay shows the number of divisions as CFSE labeling within CD3^+^ cells. Shown are all donors (K025, K029, K034) at days 4 (upper left panel), 6 (lower left panel), 8 (upper right panel) and 12 (lower right panel).

**Fig. S8:**
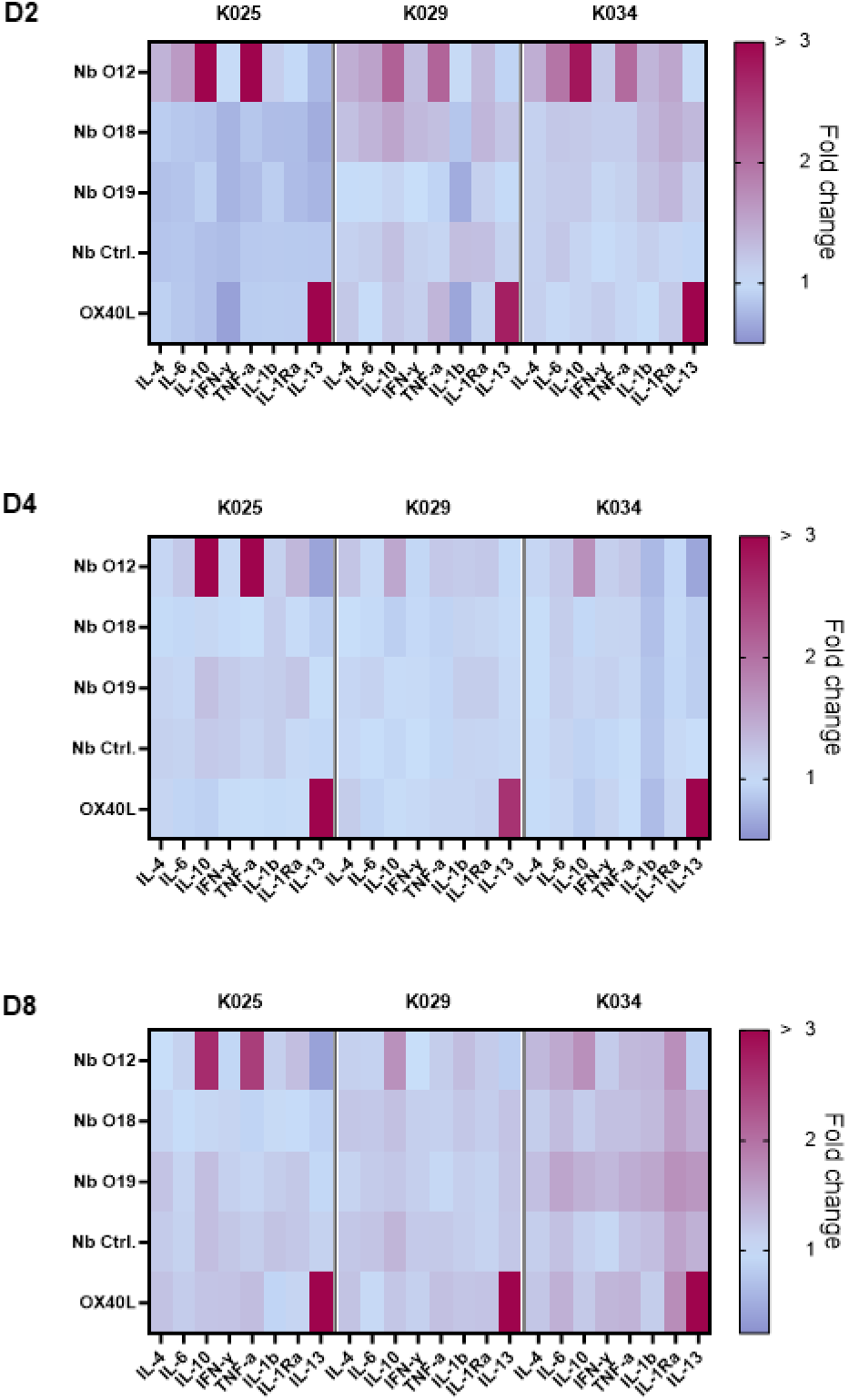
Impact of hOX40-Nb treatment on cytokine release of hPBMCs. The impact of binding of hOX40-Nbs on the release of cytokine secretion was tested by treating hPBMCs of three donors (K25, K29, K34) with 0.5 µM hOX40-Nbs, unspecific Nb (Nb Ctrl.), OX40L or left untreated upon 24h stimulation with 5 µg/mL PHA-L, to induce OX40 expression. Cytokine release to the medium was monitored at day 2, 4 and 8 (D2, D4, D8) by a microsphere-based sandwich immunoassay (see **Supplementary Table S2**) and summarized in a heat map for each time point. Values are shown as fold change compared to the untreated control based on the mean of three technical replicates (n = 3).

### Supplementary Tables

**Table S1.**
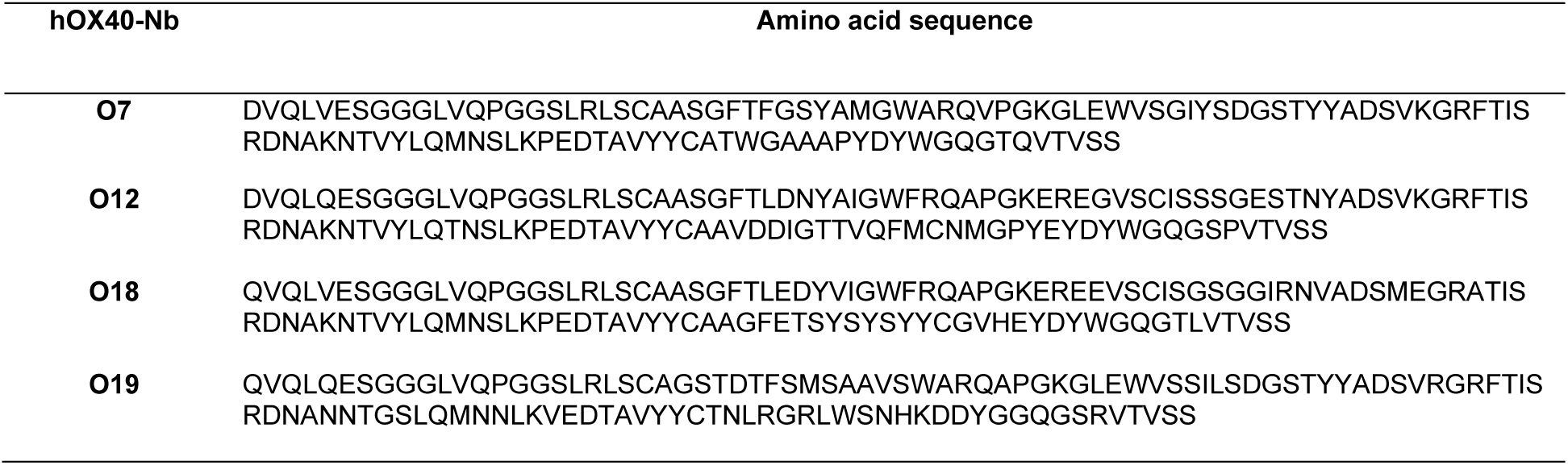
Amino acid sequences of identified hOX40-Nbs.

**Table S2.** Cytokines analyzed in this study.

| Cytokine | indicative for |
| --- | --- |
| Interleukin 1b (IL-1b) | proinflammatory |
| Interleukin 1 receptor antagonist (IL-1Ra) | antiinflammatory |
| Interleukin 4 (IL-4) | antiinflammatory |
| Interleukin 6 (IL-6) | pro-/antiinflammatory |
| Interleukin 10 (IL-10) | antiinflammatory |
| Interleukin 13 (IL-13) | antiinflammatory |
| Interferon $\gamma$ (IFN-γ) | proinflammatory |
| Tumor necrosis factor $\alpha$ (TNF-α) | proinflammatory |

**Table S3.**
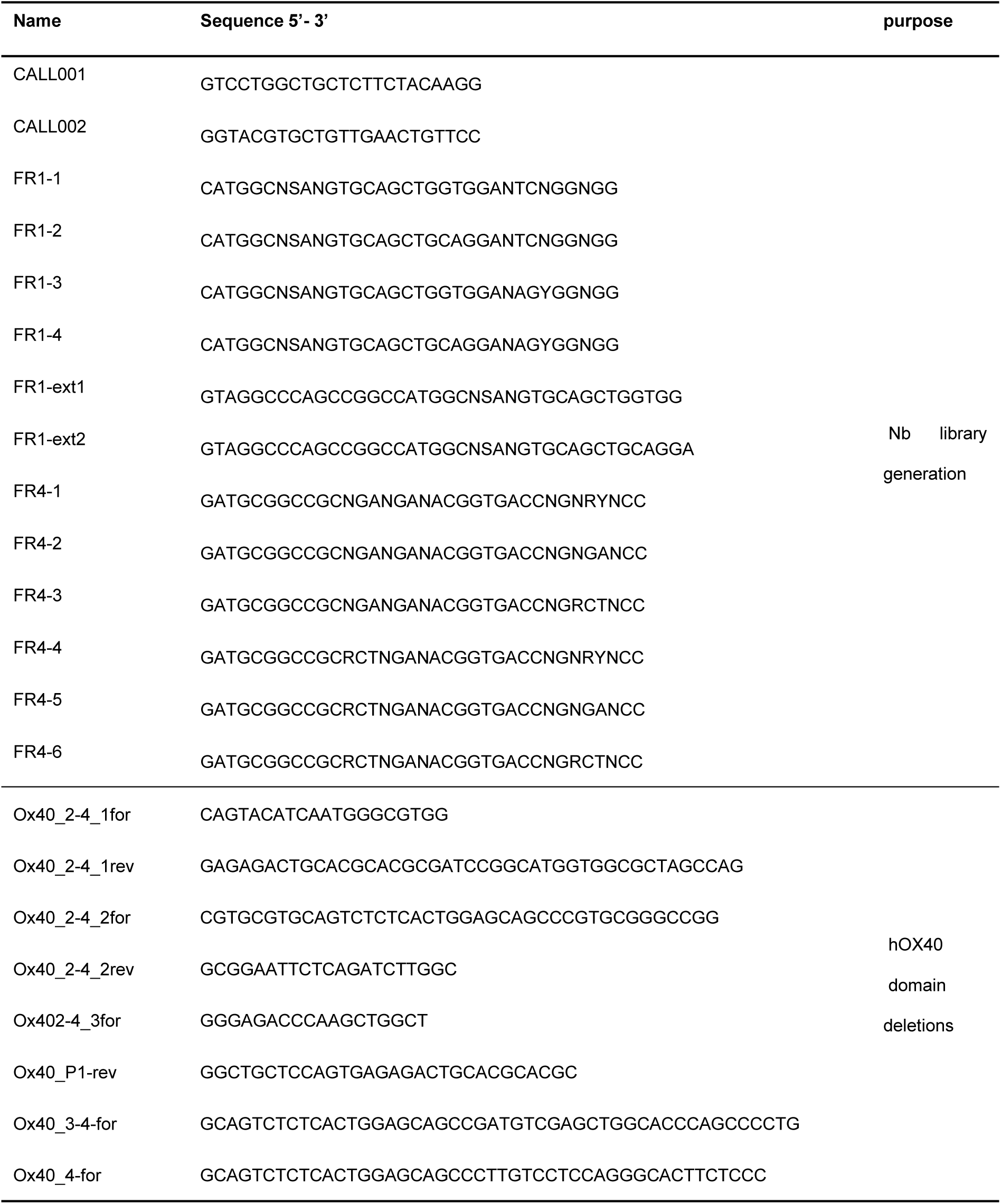
primers used in this study.

